# Non-synaptic interactions between olfactory receptor neurons, a possible key feature of odor processing in flies

**DOI:** 10.1101/2020.07.23.217216

**Authors:** Mario Pannunzi, Thomas Nowotny

## Abstract

When flies explore their environment, they encounter odors in complex, highly intermittent plumes. To navigate a plume and, for example, find food, they must solve several challenges, including reliably identifying mixtures of odorants and their intensities, and discriminating odorant mixtures emanating from a single source from odorants emitted from separate sources and just mixing in the air. Lateral inhibition in the antennal lobe is commonly understood to help solving these challenges. With a computational model of the *Drosophila* olfactory system, we analyze the utility of an alternative mechanism for solving them: Non-synaptic (“ephaptic”) interactions (NSIs) between olfactory receptor neurons that are stereotypically co-housed in the same sensilla.

We found that NSIs improve mixture ratio detection and plume structure sensing and they do so more efficiently than the traditionally considered mechanism of lateral inhibition in the antennal lobe. However, we also found that NSIs *decrease* the dynamic range of co-housed ORNs, especially when they have similar sensitivity to an odorant. These results shed light, from a functional perspective, on the role of NSIs, which are normally avoided between neurons, for instance by myelination.

**Author summary:** Myelin is important to isolate neurons and avoid disruptive electrical interference between them; it can be found in almost any neural assembly. However, there are a few exceptions to this rule and it remains unclear why. One particularly interesting case is the electrical interaction between olfactory sensory neurons co-housed in the sensilla of insects. Here, we created a computational model of the early stages of the *Drosophila* olfactory system and observed that the electrical interference between olfactory receptor neurons can be a useful trait that can help flies, and other insects, to navigate the complex plumes of odorants in their natural environment.

With the model we were able to shed new light on the trade-off of adopting this mechanism: We found that the non-synaptic interactions (NSIs) improve both the identification of the concentration ratio in mixtures of odorants and the discrimination of odorant mixtures emanating from a single source from odorants emitted from separate sources – both highly advantageous. However, they also decrease the dynamic range of the olfactory sensory neurons – a clear disadvantage.

## 1 Introduction

Flies, as most other insects, rely primarily on olfaction to find food, mates, and oviposition sites. During these search behaviors, they encounter complex plumes with highly intermittent odor signals: Odor whiffs are infrequent and odor concentration varies largely between whiffs [1–3]. To navigate a plume and successfully achieve their objectives, flies must decipher these complex odor signals which poses several challenges: Identifying odors, whether mono-molecular or a mixture; Identifying odor intensity; Discriminating odorant mixtures emanating from a single source from those emanating from separate sources; identifying source locations, etc. Early sensory processing is understood to play an important role for solving these challenges [4–6]. For instance, lateral inhibition in the antennal lobe is commonly understood to be useful for decorrelating odor signals from co-activated receptor types. Here we investigate the hypothesis that the early interactions between ORNs in the sensilla are similarly, if not more, useful for decoding information in odor plumes.

In both, vertebrates and invertebrates, odors are sensed by an array of numerous receptor neurons, each typically expressing receptors of exactly one of a large family of olfactory receptor (OR) types. In *Drosophila*, olfactory receptor neurons (ORNs) are housed in evaginated sensilla localized on the antennae and maxillary palps [7], each sensillum containing one to four ORNs of different types [7, 8]. The co-location of ORN types within the sensilla is stereotypical, i.e. ORNs of a given type “a” are always co-housed with ORNs of a specific type “b” [8]. Furthermore, ORNs within the same sensillum can interact [9–13] without making synaptic connections (see Fig.1a). While the interactions are sometimes called “ephaptic”, referring to their possible electric nature [14], we here prefer to call them non-synaptic interactions (NSIs), for the sake of generality. Whether stereotypical co-location of – and NSIs between – ORNs have functions in olfactory processing and what these functions might be remains unknown, even though several non-exclusive hypotheses have been formulated (see Discussion and [6, 11, 14] and references therein).

**Fig 1.**
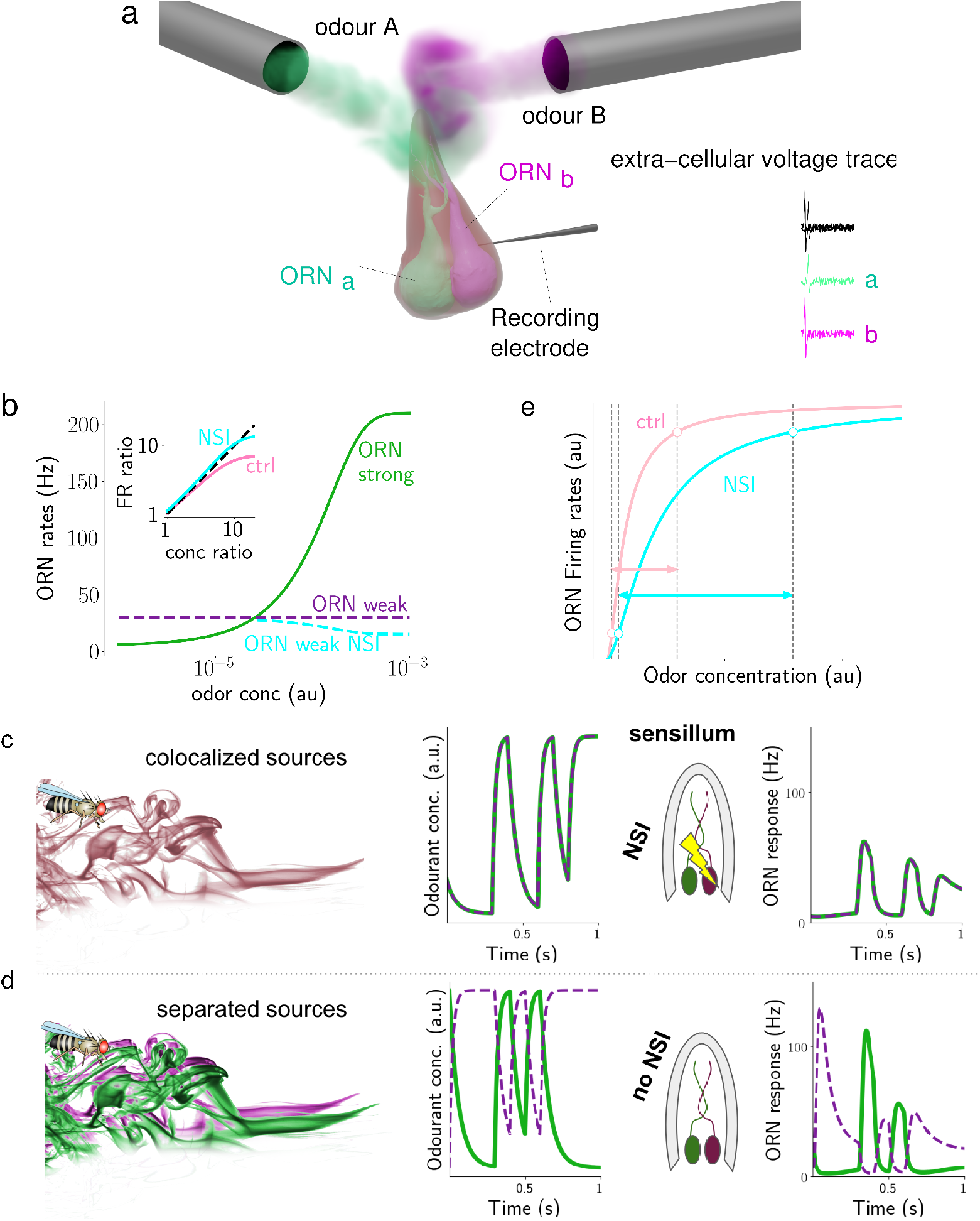
a) Non-synaptic interaction (NSI). Theoretical and experimental studies have proposed that the NSI between ORNs is mediated by a direct electrical field interaction between such closely apposed neurons. b) **Hypothesis 1**: Due to the sigmoidal shape of the dose-response curve (green line), the concentration ratio is not well encoded (pink line in inset). The second ORN’s activity is indicated with a purple dashed line. When NSIs are present, the second ORN (cyan dashed line) is inhibited by the first more strongly activated ORN, which improves the ratio encoding (cyan line in the inset), making the encoding ratio more similar to the concentration ratio (black dashed line). **Hypothesis 2**: If a single source emits an odorant mixture (c), odorants will arrive in close synchronization, NSIs will take effect and the response in both ORNs is affected. If separate sources emit the odorants (d), they will arrive in a less correlated way [15], and NSIs have almost no effect, resulting in larger ORN responses. ORN response data shown is based on a preliminary model. **Hypothesis 3**: The lateral inhibition caused by the presence of NSIs decrease the ceiling effect for high concentration and therefore shift the right boundary of the dynamic range to higher concentrations, thus increasing the dynamic range (panel e).

Here, we investigate three hypotheses: First, NSIs could help the olfactory system to identify ratios of odorant concentrations in mixtures more faithfully by enhancing the dynamic range over which ORN responses contain the relevant information to do so (see Fig.1, panel b). Second, NSIs could help improve the spatiotemporal resolution of odor recognition in complex plumes (see Fig.1, panels c-d). Third, NSIs could improve the dynamic range of ORNs’ responses to individual odorants by partially removing the ceiling effect that occurs for high concentrations. In all three hypotheses, the NSI mechanism has to compete with lateral inhibition in the antennal lobe, which is commonly recognized to fulfill these roles, even though, of course, the two mechanisms are not mutually exclusive.

Indirect support for the first hypothesis is found in the context of moths’ pheromone communication. In some moth species, pheromone mixture ratio discrimination is critical for survival and therefore even slight changes in pheromone component ratios of 1-3% can cause significant changes in behavior. In these species, the ORNs responding to pheromone components are more likely to be co-housed. Meanwhile, when mixture ratios are not as critical for behavior, i.e., significant changes in behavior only occur if pheromone component ratios change 10% or more, ORNs are less likely to be paired in the same sensilla (see [11] and references therein). The problem when encoding ratios of odor concentration with activity of PNs (or ORNs) is that, while concentration spans several orders of magnitude, their firing rates can only span 2 orders of magnitude. Stimulus intensities in receptor neurons are commonly considered to be coded by spiking frequency (at least since [16]). A concentration ratio around 10 can be difficult, or impossible, to encode if the weakly activated ORN’s activity is already around 50 Hz. NSIs would decrease the activity of the weakly activated ORN, inhibited by the activity of the more strongly activated ORN. In this way, the encoding of the ratio can be improved (see Fig.1b).

The improvement of spatiotemporal resolution of the second hypothesis can be achieved by decorrelating odor response profiles to improve odor recognition (see Fig.1, panels c-d), again much like lateral inhibition in the antennal lobe (AL), or centre-surround inhibition in the retina. Odorants dissipate in the environment in complex, turbulent plumes of thin filaments of a wide range of concentrations, intermixed with clean air. Odorants emanating from the same source presumably travel together in the same filaments while odorants from separate sources are in separate strands (see e.g., [15] for empirical evidence for this intuitive idea). Insects are able to resolve odorants in a blend and recognize whether odorants are present in a plume and whether or not they belong to the same filaments [17–21]. In the pheromone sub-system of moths, it is known that animals are able to detect, based on fine plume structure, whether multiple odorants have been emitted from the same source or not [17, 18, 22]. In the pheromone subsystem of *Drosophila*, ORNs responding to chemicals emitted by virgin females and ORNs responding to chemicals emitted by mated females are co-housed in the same sensilla: The ‘virgin females ORNs’ on the males’ antenna promote approach behavior, but the ‘mated females ORNs’ inhibit ‘virgin females ORNs’ [23]. This inhibition could be implemented through NSIs [8, 11, 23, 24].

The third hypothesis assumes that NSIs can affect the dynamic range and the odor detection threshold. It was motivated by the work of Vermeulen and Rospars [14] who first modelled the ORNs’ interactions with an electrical circuit, hypothesizing that electrical insulation of the sensillar lymph from the hemolymph generates a common potential difference between the dendritic and axonal compartments of the ORNs within a sensillum. As a consequence, when an odorant activates an ORN, the resistance of the insulation would drop and the activity of the co-localized ORN would show a reduced receptor potential. At the same time, the receptor potential of the activated neuron would be increased, leading to a stronger response of the more sensitive ORN and a slightly lower odor detection threshold. These effects are strongest if the sensitivities to the analysed odorant are different. The model has been referred to as evidence for the benefits of co-housed ORNs with NSIs throughout the literature [8, 10, 13, 25, 26]. Here, we analyse the effects of NSIs for an individual odorant beyond the previously considered steady state equations and include the known spike rate adaptation of ORNs (see e.g. [27, 28]) as well.

The experimental evidence for these hypotheses and for the general relevance of NSIs for olfactory processing remains mixed and research is still at an early stage (but see [6] for a more detailed analysis). Encouraged by the available evidence, and without trying to rule out other hypotheses (for further analysis see Discussion), our goal is to investigate, with a computational model, the viability of the hypothesized functions of NSIs between ORNs. Our computational approach helps experimental studies to refine hypotheses about NSIs and eventually answer the pertinent question why such a mechanism that appears to duplicate what is already known to be implemented by local neurons in the AL could nevertheless provide an evolutionary advantage.

A number of computational models have been developed to capture different aspects of the olfactory system of insects. However, until recently, most modeling efforts were based on the assumption of continuous constant stimuli, which are partially realistic only for non-turbulent fluid dynamics regimes (see [29], and reference therein). Most commonly insects encounter turbulent regimes, in which odorant concentration fluctuates rapidly (see SUPP S5).

To cope with these more realistic stimuli, [27, 30–32] have formulated new models of *Drosophila* ORNs, that are constrained by experimental data obtained with more rich, dynamic odor inputs, including a model simulating ORNs and PNs that are subject to input from simulated plumes [32] with statistical properties akin to those of naturalistic plumes (see more details in Model and methods and Correlation detection in long realistic plumes).

Here, we present a network model with two groups of ORNs, each tuned to a specific set of odorants, connected to their corresponding glomeruli, formed by lateral neurons (LNs) and PNs, following the path started by [33, 34], and subsequently by [35–37]. We model the ORNs in a similar approach as [27, 30] with minor differences in the filter properties and the adaptation (see Model and methods). We have tested the behavior of this network in response to simple reductionist stimuli (as commonly used in the literature, see above), and simulated naturalistic mixtures plumes (as described by the experiments in [2, 3]). We then used this simple but well-supported model to investigate the role of NSIs for odor mixture recognition.

## 2 Results

To investigate the role of NSIs in olfactory sensilla, we have built a computational model of the first two processing stages of the *Drosophila* olfactory system. In the first stage, ORN responses are described by an odor transduction process and a spike generator (see Model and methods), in line with previous experimental and theoretical studies [27, 28, 30, 38]. We simulated pairs of ORNs expressing different olfactory receptor (OR) types, as they are co-housed in sensilla. NSIs between co-housed pairs effectively lead to their mutual inhibition (see Fig.1a). The second stage of olfactory processing occurs in the AL, in which PNs receive input from ORNs and form local circuits through LNs. ORNs of the same type all make excitatory synapses onto the same associated PNs. PNs excite LNs which then inhibit PNs of other glomeruli but not the PNs in the same glomerulus (see Fig.2 and Model and methods for further details).

**Fig 2.**
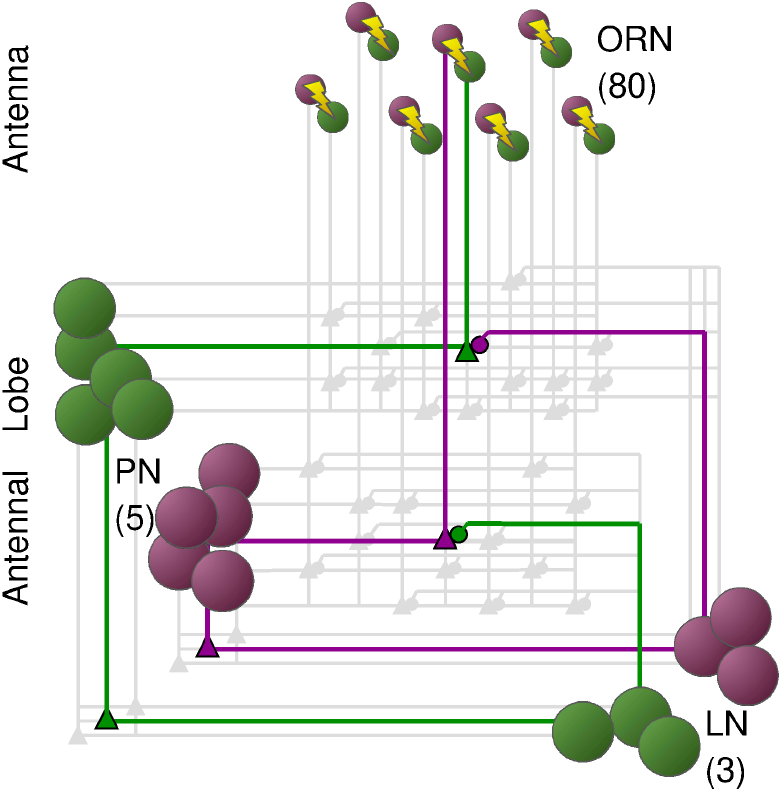
Model of the early stages of the olfactory system of the *Drosophila*. The model simulates the odor transduction, the ORNs and the AL using only two groups of ORNs (ORN_*a*_ and ORN_*b*_) and their respective PNs and LNs. Each ORN type, a and b, is tuned to a specific set of odorants (e.g. individual pheromone component) and converges onto its corresponding PNs. PNs impinge into their respective LNs, but receive inhibitory input from LNs of the other type.

For maximum clarity, we here focus on only one type of sensillum and hence two types of ORNs that we denote as ORN_*a*_ and ORN_*b*_. We initially assume that odorants labeled A and B selectively activate ORN_*a*_ and ORN_*b*_, respectively (see Fig.2 and Fig.1a), but we relax this constraint when analyzing the effects on the ORNs’ dynamic range. This assumption is not only sensible for a reductionist analysis of the role of NSIs, but it is also based on experimental observations. For instance, pheromone receptors in moths and in *Drosophila* are highly selective, paired in sensilla, and exhibit NSIs [11, 39]. In the general olfactory system of *Drosophila*, neurons ab3A and ab3B in sensillum ab3 are selectively sensitive to 2-heptanone and Methyl hexanoate, and when stimulated simultaneously they inhibit each other through NSIs [10].

### 2.1 Constraining the ORN model to biophysical evidence

In this investigation we are particularly interested in the complex time course of odorant responses and have therefore focused on replicating realistic temporal dynamics of the response of ORNs at multiple time scales. ORN responses were constrained with experimental data obtained with delta inputs, i.e. inputs of very short duration and very high concentration, and random pulses, i.e. series of input pulses which durations and inter-stimulus-intervals were drawn from distributions (see SUPP Fig.S5) extrapolated from recordings of odorant activity in open space [1–3]. We found that our model reproduces the data to a similar quality (percentage error of around 10% for high activity, >SI30Hz) as previous linear-nonlinear models [27, 28, 30, 38, 40], even though it has fewer free parameters (see Fig.3).

**Fig 3.**
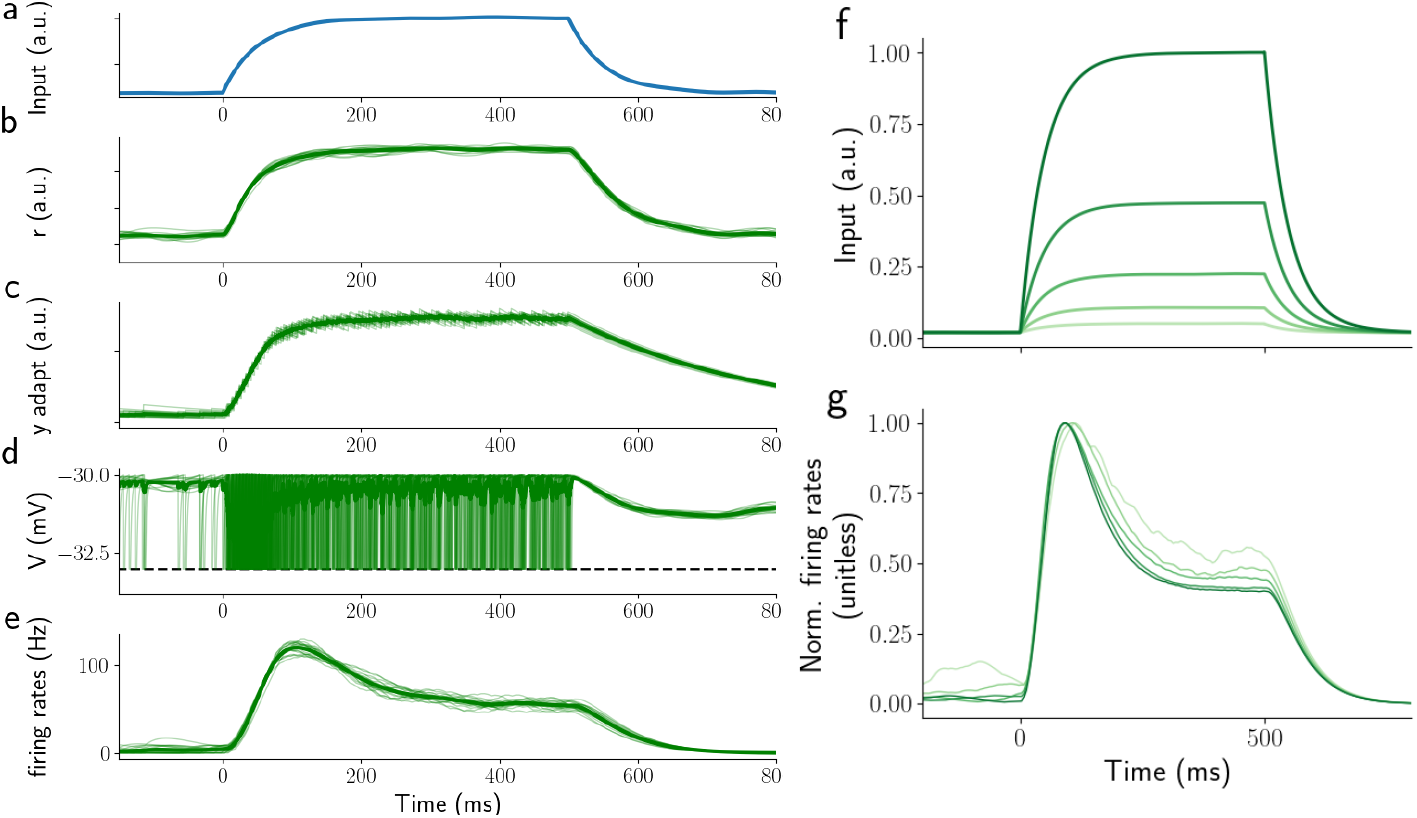
ORN responses to a 500 ms single step stimulus. Panel a) shows stimulus waveform, panel b) response of the activation variable *r* of the transduction phase, panel c) activity of the internal ORN variable y (see Model and methods), panel d) ORN potential response of all ORNs, panel e) Spike density function of the ORN population activity. Panel f) stimulus waveforms for different odorant concentrations, panel g) ORN activity normalized to the peak activity. Odor concentration is indicated with different shades of green. After normalization, the responses are almost identical to those reported by [28].

To further constrain the model, we compared its results to electrophysiological recordings from ORNs [27, 38] responding to 2 s long odor stimuli with shapes resembling steps, ramps, and parabolas (see SUPP Fig.S2 and Model and methods). The model reproduces all key properties of the experimentally observed ORN responses. For the step stimuli, ORN activity peaks around 50ms after stimulus onset and the peak amplitude correlates with the odor concentration (SUPP Fig.S1a-b). After the peak, responses gradually decrease to a plateau. Furthermore, if normalised by the peak value, responses have the same shape independently of the intensity of the stimulus [28], see Fig.3f,g. For the ramp stimuli, ORN responses plateau after an initial period of around 200 ms, encoding the steepness of the ramp (see SUPP Fig.S1c-d). More generally, ORN responses seem to encode the rate of change of the stimulus concentration [27, 38, 40]. Accordingly, ORN activity in response to the parabolic stimuli is like a ramp (see SUPP Fig.S2e-f). Parameters used for the transduction and the ORN dynamics are reported in Table 1.

**Table 1.**
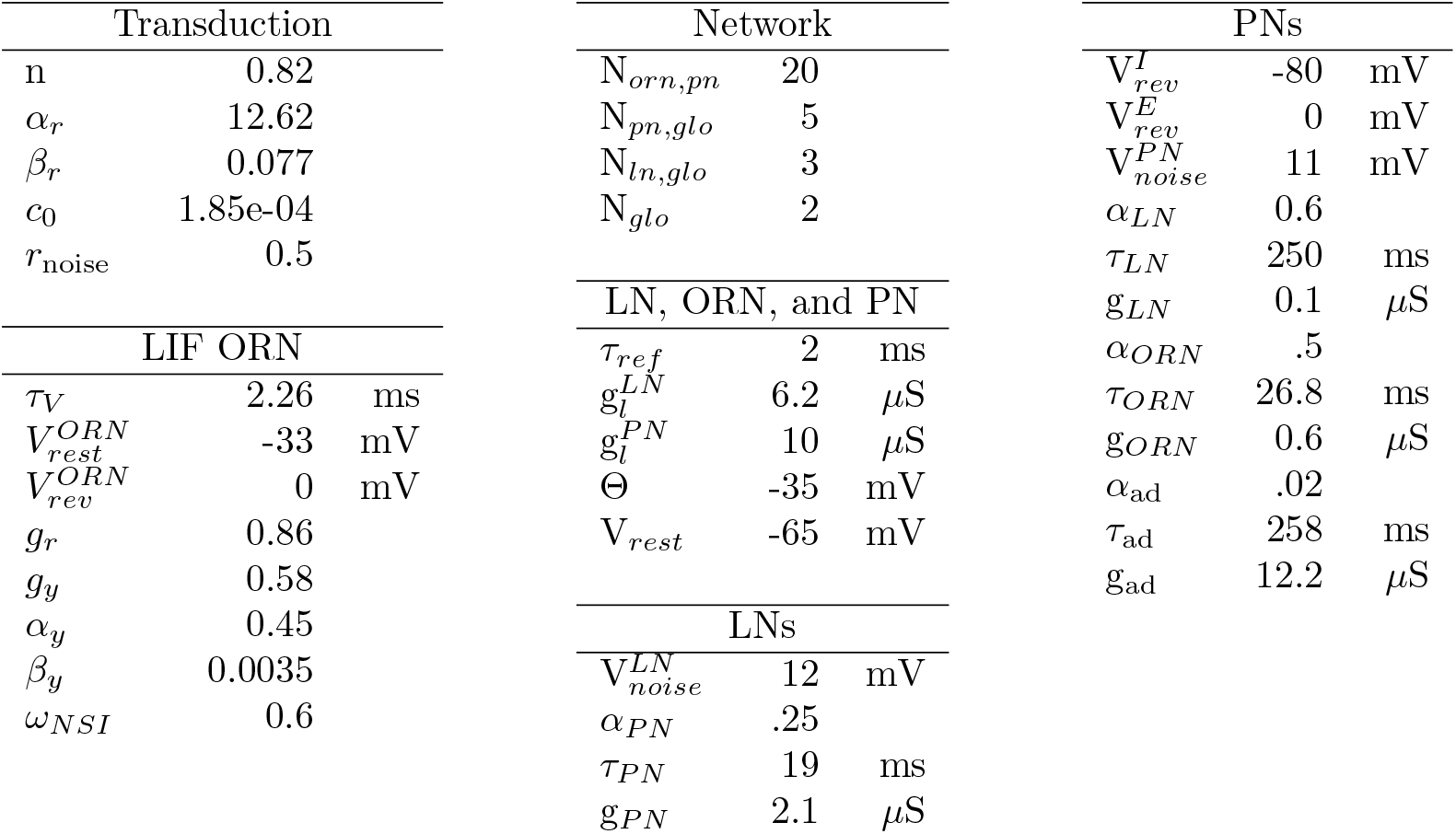
Model parameters. To reproduce the experimental data in our model, we used the following parameters: Transduction, LIF ORNs, PNs and LNs, and Network parameters. We fitted a subset (24) of ORN, PN and LN parameters in order to reproduce the time course shown in e-phys experiments (e.g. [27, 28]); we obtain similar correlated values as those reported in [82] by fitting the amount of noise injected in the 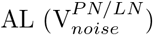 and in the transduction layer (c_0_); Network parameters are not fitted, but extracted from the literature (e.g. [82, 88, 89]). NSI strength (*w*_*NSI*_) and synaptic strength of LNs (*α*_*LN*_) are not fitted, but their values were changed to explore the network behavior.

Finally, we fitted the parameters of PNs so that the PN responses with a single constant step stimulus resemble the qualitative behavior reported in [41]. In that study, the authors reported that the response of PNs to such a stimulus is best described by a sigmoid,

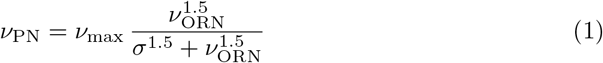

where *ν*_max_ is the maximum firing rate of ORNs, *σ* is a fitted constant representing the level of ORN input that drives half-maximum response, and *ν*_ORN_ and *ν*_PN_, are the average firing rates of the ORNs and the PN over the stimulation period (500 ms), respectively. Our model is able to reproduce this behavior (see 4e). LNs follow the same behavior (see Fig.4f). Note that this result, i.e. the sigmoidal behavior, generalizes to both, shorter stimulation times (50 and 100 ms, see SUPP Fig.S3 and Fig.S4) and to the peak activity instead of the time averaged activity. See Table 1 for the remaining parameters of the model.

**Fig 4.**
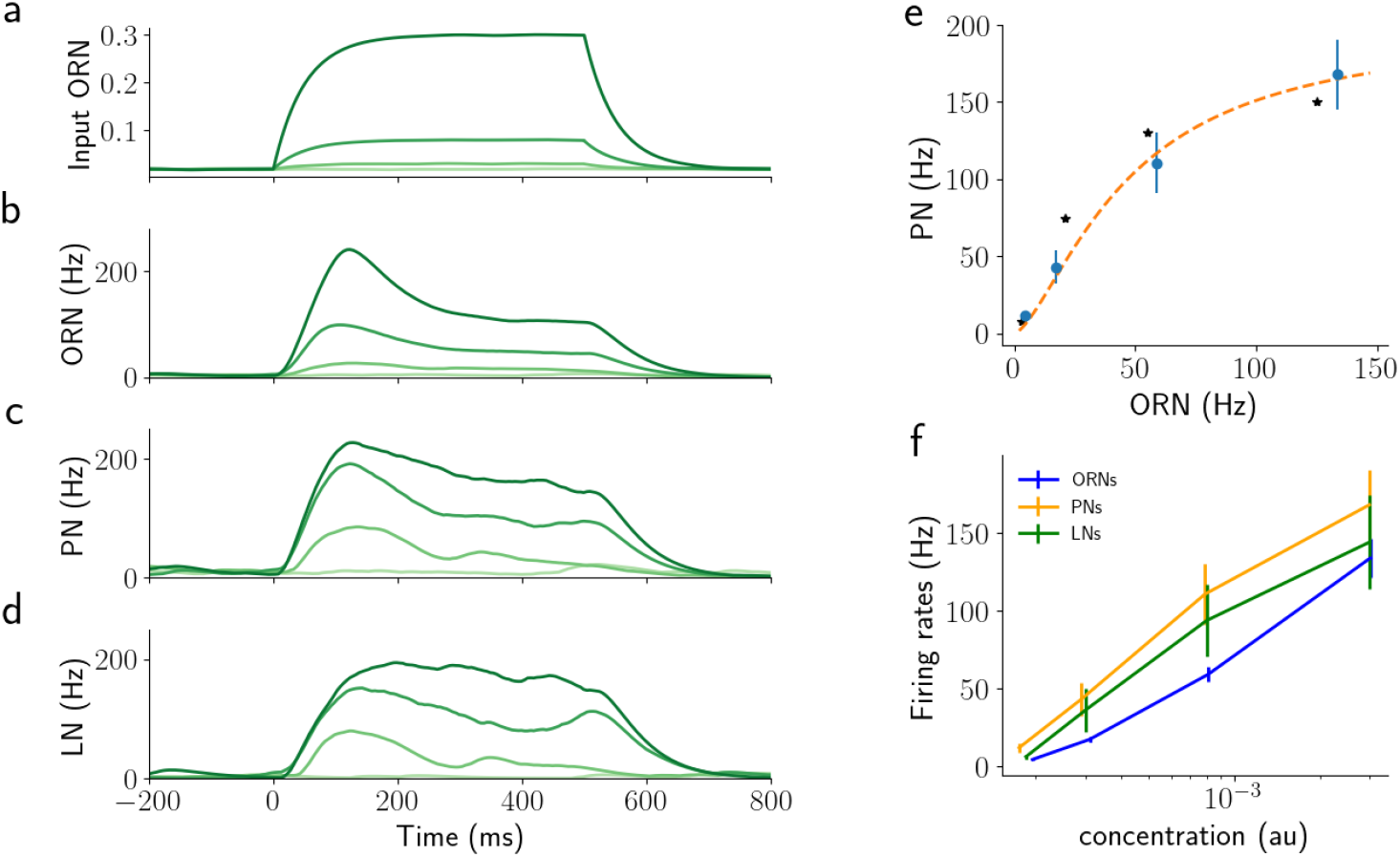
Network response to 500 ms step stimuli of a single odorant for the network as shown in 2. panel a) shows step stimuli, shade of green indicates concentration, panels b-d) corresponding activity of ORNs, PNs, and LNs. Shades of green match the input concentrations. e) Average response of PNs over 500 ms against the average activity of the corresponding ORNs. The orange dashed line is the fit of the simulated data using equation eq. 1 as reported in [41]. Asterisks are the values reported in [41]. f) Average values for PNs, ORNs, and LNs for different values of concentration. Error bars show the SE over PNs.

With a model in place that demonstrates the correct response dynamics for a variety of stimuli, we then analysed its predictions on whether NSIs can be beneficial for odor mixture processing. In particular, we tested the following three hypotheses: 1. NSIs improve the encoding of concentration ratio in an odorant mixture (see section Odorant ratio in synchronous mixture stimuli), 2. NSIs support differentiating mixture plumes from multiple versus single source scenarios (see section Processing asynchronous odor mixtures), NSIs increase ORNs dynamic range (see section Increasing ORN Dynamic Range and sensitivity). For the first two hypotheses, we compared the model with NSIs (“NSI model”) to a model with lateral inhibition between PNs mediated by LNs in the AL (“LN model”) and a “control model” where the pathways for different OR types do not interact at all (see Fig.6).

**Fig 5.**
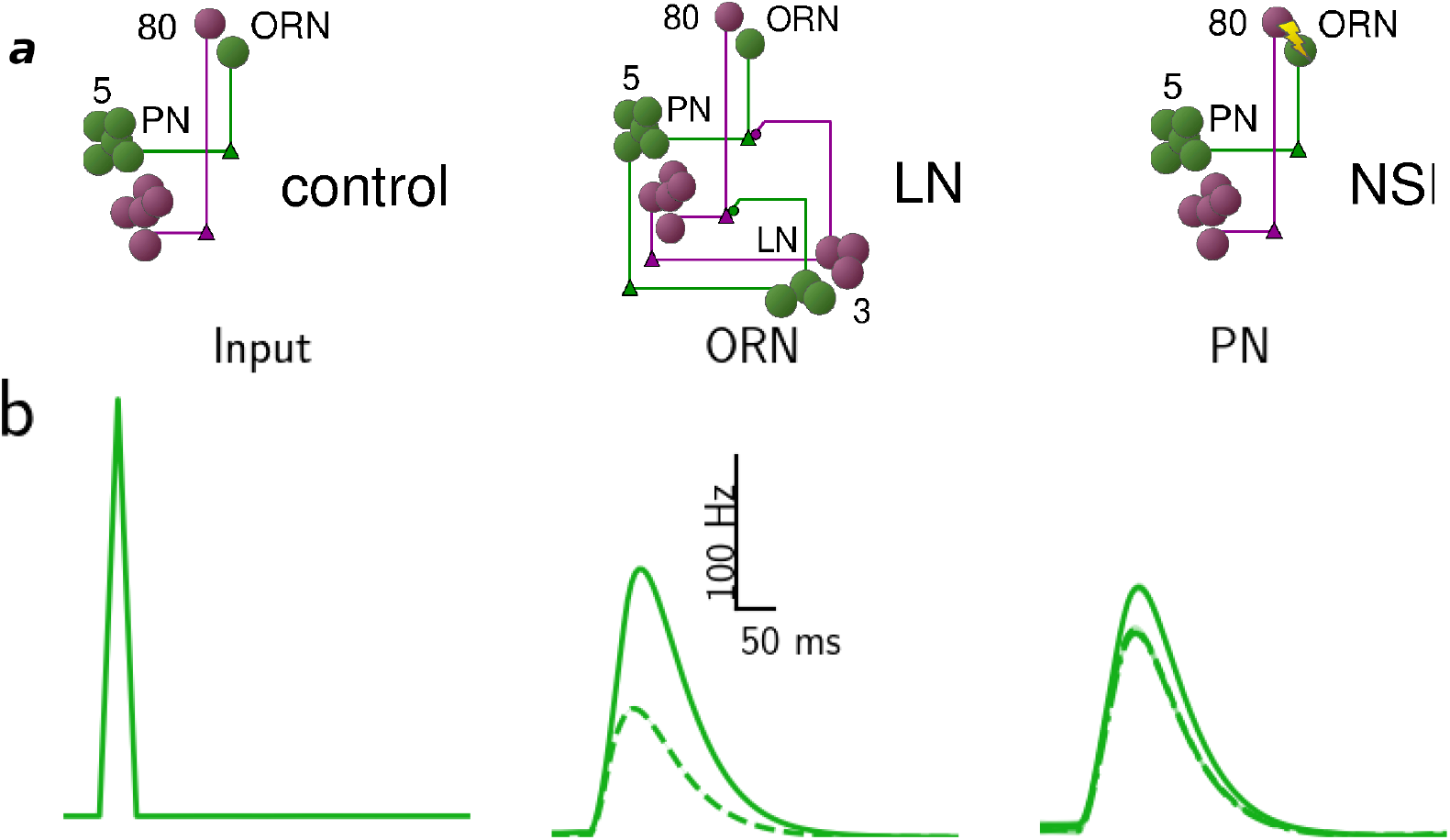
Responses to synchronous pulses of the control, LN and NSI model. Panel a) Schematics of the three models analyzed: control, LN and NSI. Panel b) shows the time course of ORN and PN activity in response to a triangular pulse (50 ms, 1st column) for the three models – control (continuous line), NSI (dashed line) and LN model (dot and dashed line). Input to the three models is identical, while control and LN models have identical ORN activity, which is therefore not displayed twice. The lines show the average response and the shaded area around the lines the standard deviation over 10 trials.

**Fig 6.**
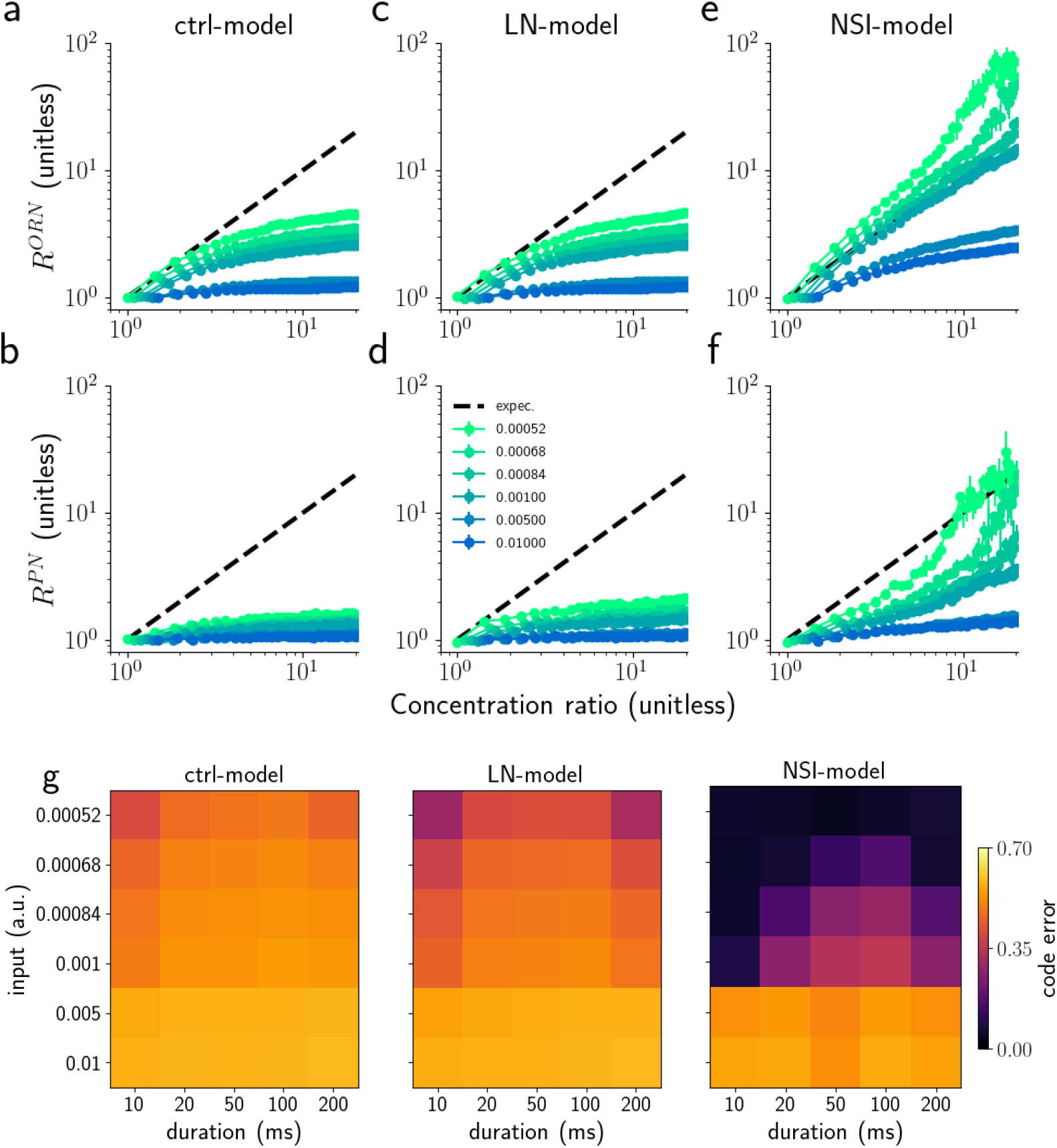
Encoding concentration ratio with average PN activity. ORN (a,c,e) and PN (b,d,f) responses ratio (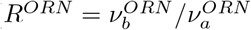 and 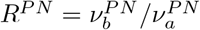 to a single synchronous triangular pulse of 50 ms duration applied to both ORN groups. The graphs show response ratios versus concentration ratio of the two odorants for four different overall concentrations (colours, see legend in f). The peak PN responses would be a perfect reflection of the odorant concentration if they followed the black dashed diagonal for all concentrations. Error bars represent the semi inter-quartile range calculated over 10 trials. g) Analysis of the relative distance (that is a coding error) for different values of stimulus duration and concentration. The coding error is calculated as the squared relative distance (see Model and methods).

### 2.2 Odorant ratio in synchronous mixture stimuli

Airborne odors travel in complex plumes characterized by highly intermittent whiffs and highly variable odor concentration [1–3]. To successfully navigate such plumes and find for example food, flies must recognize relevant whiffs regardless of the overall odor concentration in them, i.e. perform concentration-independent odorant ratio recognition. This is a difficult problem mainly because PN responses are sigmoidal with respect to concentration. To investigate this problem and understand whether NSIs may play a role in solving it, we stimulated the model ORNs with binary mixtures, varying the overall concentration of the mixture, the concentration ratio, and the onset time of the two odorants. The whiffs in plumes (see e.g. SUPP Fig.S5a) are mimicked with simple triangular odorant concentration profiles that have a symmetric linear increase and decrease (see Fig.5a). We first analysed the case of synchronous pulses, which is typical when a single source emits both odorants (see SUPP Fig.S8 a, extracted from [15]).

Fig.5b shows the typical effects of the NSI mechanism and AL lateral inhibition on PN responses. For the purpose of this figure we adjusted the NSI strength and LN synaptic conductance in such a way that the average PN responses to a synchronous mixture pulse were similar across the two models (see Table 1). While the stimulus lasts only 50 ms, the effect on ORNs, PNs and LNs lasts more than twice as long. We observed the same behavior for other stimulus durations (tested from 10 to 500 ms). In the control model (continuous lines), PN and LN responses are unaffected by lateral interactions between OR-type specific pathways and because we have matched the sensory response strength of the two odorants and OR types for simplicity, the responses of the PN in the two glomeruli are very much the same. For the LN model (dot and dash lines) the response of ORNs is unaltered by network effects and synaptic inhibition of LNs is the only lateral interaction between pathways. For the NSI model (dashed lines), ORN activity is directly affected by NSIs and the activity of PNs is lower than in control conditions as a consequence of the lower ORN responses. As explained above, NSI strength and synaptic conductance of LN inhibition were chosen in this example so that the response of the PNs for both models is of similar magnitude.

#### 2.2.1 Encoding concentration ratio for synchronous stimuli

To investigate the effectiveness of the two mechanisms for ensuring faithful odorant ratio encoding more systematically, we tested the three models with synchronous triangular odor pulses of different overall concentration, different concentration ratios, and for different values of stimulus duration (from 10 to 200 ms), which we selected to match the range of common whiff durations observed experimentally (see SUPP Fig.S5). Here, and throughout the study we explored several values for the two strength parameters (0.1 to 0.6 for both *ω*_*NSI*_ and *α*_*LN*_) and for each analysed task we report the results of the best performing NSI and LN model, respectively. The results are summarised in Fig.6 based on the ratio of peak activity *R* = *ν*_*b*_*/ν*_*a*_ (both for ORNs and PNs, see Model and methods) during the first 200 ms after the stimulus onset.

Due to the fundamentally sigmoidal relationship between ORNs and PNs responses and odorant concentration (see Fig.4e), the encoding of the ratio between two odorants in a mixture is distorted (see the control model response in Fig.6a,b). The encoding of odorant mixtures is indeed already disrupted at the level of the ORNs (Fig.6a), not only on the level of PNs (Fig.6b). Once activated, inhibitory interactions between PNs mediated by LNs slightly improve ratio encoding (Fig.6c,d), reducing the strong ceiling effect for high concentration. The NSI mechanism instead clearly changes the ORN responses (Fig.6e), and as a consequence, PN responses change so that their activity reflects the ratio of odor concentrations better for most of the tested concentration ratios (Fig.6f).

The same results shown for peak activity are true for the average activity calculated during the first 200 ms after the stimulus onset (see SUPP Fig.S6). In the same vein, testing with longer stimuli also yielded qualitatively similar results.

Next, we tested the encoding of ratios – besides for different concentrations – also for different whiff durations. In Fig.6g we show the coding error (here measured as relative distance of the concentration ratio and PN ratio) for different values of stimulus duration and concentration. It is clear that NSI model outperforms both LN and control model, apart for very high concentrations. While very intuitive, encoding mixture ratios linearly in PN firing rates is not the only option. To analyse encoding quality more generally, we therefore repeated our analysis by calculating the mutual information (MI) between the odorant concentration and *R*^*PN*^. The results from the analysis of the MI are qualitatively similar to the results with the coding error (see SUPP Fig.S7).

In the next section we will explore the effectiveness of the different models when the whiffs arrive asynchronously.

### 2.3 Processing asynchronous odor mixtures

When odorants are released from separate sources, they form a plume in which the whiffs of different odorants typically are encountered at distinct onset times. To the contrary, when a mixture of odorants is released from a single source they form a plume where the odorants typically arrive together (SUPP Fig.S8). We hypothesise that if lateral inhibition (via LNs or NSIs) only takes effect in the synchronous case but not in asynchronous case, it will help distinguishing single source and multi-source plumes. For instance, in the case of pheromone receptor neurons that are co-housed with receptor neurons for an antagonist odorant, the response to the pheromone would be suppressed by NSI when both odorants arrive in synchrony (same source) and not when arriving with delays (the pheromone source is separate from the antagonistic source). This is thought to underlie the ability of male moths to identify a compatible female, where the antagonist odorant is a component of the pheromone of a related but incompatible species [18]. To test whether this idea is consistent with the effect of NSIs as described by our model, we calculated the predicted responses of PNs to asynchronous whiffs of two odorants in our three models - *control*, with *LN inhibition*, and with *NSI*.

Fig.7a shows the responses in the models for the example of two 50 ms triangular odor pulses of the same amplitude and at 50 ms delay. We chose stimuli that excite the two ORN types with the same strength to simplify the analysis and focus on the differences between models with respect to asynchronous input rather than differing input ratios that we analyzed above. In the control model, responses are very similar between ORN_*a*_ and ORN_*b*_, as well as, PN_*a*_ and PN_*b*_ as expected in the absence of interactions and for identical stimulus strength.

**Fig 7.**
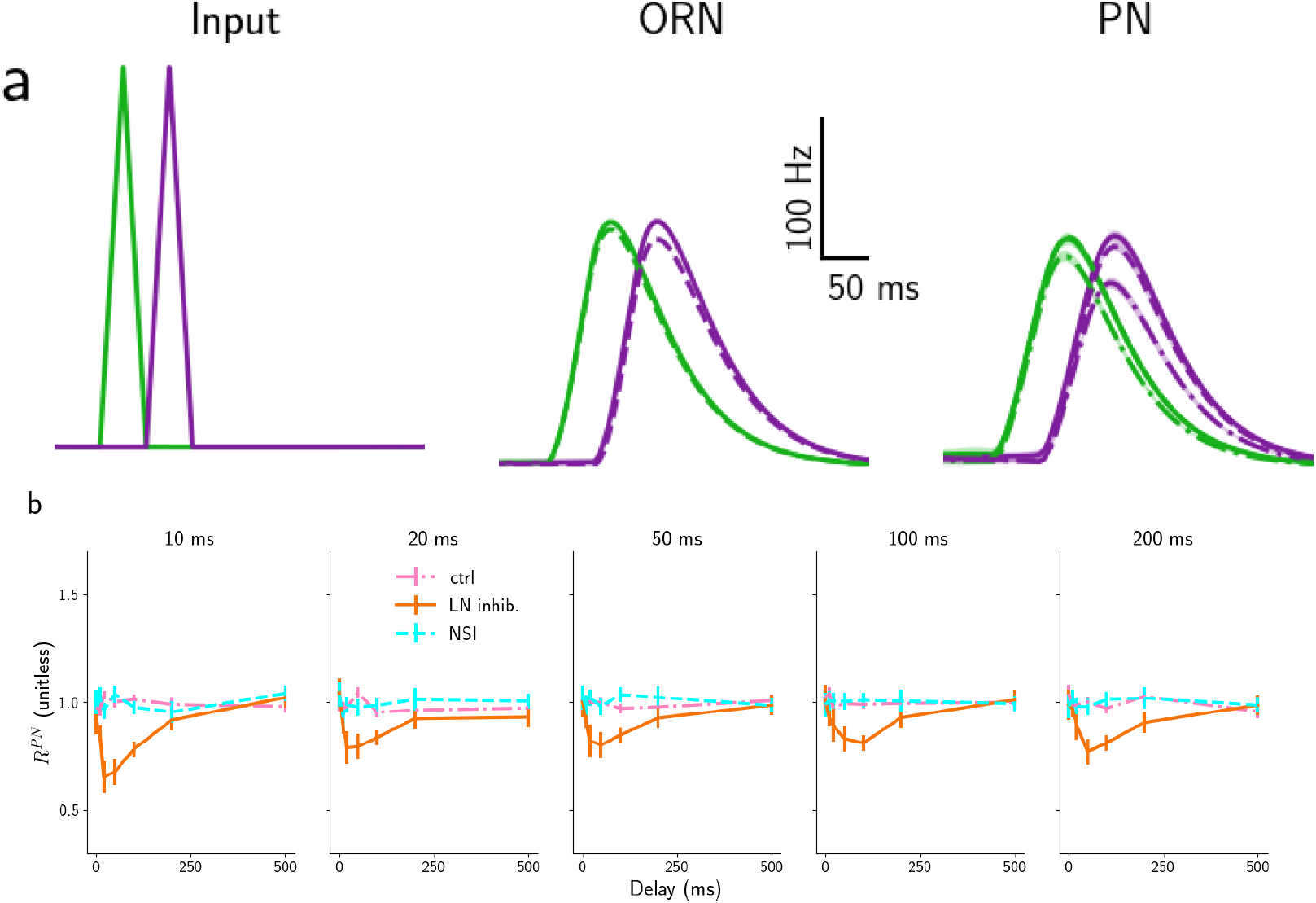
Responses to asynchronous pulses of the control, LN and NSI model. Panel a) Time course of ORN and PN activity in response to two asynchronous triangular pulses (duration 50 ms and delay 50 ms) for the three models – control (continuous line), NSI (dashed line) and LN model (dot and dashed line). Input to the three models is identical, while control and LN models have identical ORN activity, which is therefore not displayed twice. The two colours represent the two odors, ORN and PN types. The lines show the average response and the shaded area around the lines the standard deviation over 10 trials. Panel b) Median ratio of the peak PN responses of the two glomeruli 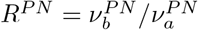 in the three models: control model (dot dashed pink), LN model (orange continuous), and NSI model (dashed cyan) for different stimulus durations as marked on the top. Error bars represent the semi inter-quartile ranges. The same plots are shown for *τ*_*LN*_ is low (25 ms) in SUPP Fig.S10.

The situation is different when LN inhibition is present (dot and dashed lines). Even for the comparatively large delay of 50 ms – the second stimulus starts when the first one ends – for the *LN model* (dot and dashed line) the excitatory input to PN_*b*_ (purple) is diminished by the inhibition coming through the LNs activated by PN_*a*_ (green). This is a consequence of PN and LN responses outlasting the stimuli as observed above. While no inhibitory effect is present in the NSI model (dashed lines), or it is much weaker. This difference depends on its characteristic time (*τ*_*LN*_). For short *τ*_*LN*_ (25 ms), the behavior of the *LN model* is almost indistinguishable from the *NSI model* (see Fig.**??**a).

To quantify the differences between the three models across different typical conditions, we calculated the ratio between the PN responses of the two glomeruli 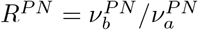, both for the peak activity and for the average activity over the stimulation time. Fig.7b shows the results for stimulus durations between 10 and 200 ms and delays from 0 to 500 ms. The whiff durations and delays were selected to match the range of values commonly observed in experiments (see SUPP Fig.S5).

As expected, the value of *R*^*P N*^ is close to 1 (pink lines) for the control model with independent ORNs and PNs and all explored parameters. In contrast, the LN model exhibits clear effects of their lateral interactions: The response of the second PN_*b*_ is suppressed by the response to the first stimulus for all tested whiff durations and for all delays shorter than 500 ms. The NSI model shows no suppression, but it is not completely without interference problems: for short delays, when there is an overlap between the two stimuli, and for very high odorant concentrations, the response of the first ORN_*a*_ is more inhibited by the activation of the second ORN_*b*_, than the second by the first. However, this effect disappears for non-overlapping stimuli.

The results are very similar whether measured in terms of the peak activity (Fig.7) or the average activity over the stimulus duration. The overall effect on the LN model is only slightly diminished for short *τ*_*LN*_ (25 ms, see SUPP Fig.S10).

This analysis clearly shows another advantage of the NSI over the LN model.

### 2.4 Correlation detection in long realistic plumes

So far we have seen that NSIs are beneficial for ratio coding in synchronous mixtures and that they distort responses less than LN inhibition in the case of asynchronous mixtures. In this section, we investigated and compared the effects of the two mechanisms when the system is stimulated with more realistic signals of fluctuating concentrations that have statistical features resembling odor plumes in an open field (see SUPP Fig.S5). The statistics of the plumes and how we simulated them are described in detail in the Model and methods; in brief, we replicated the statistical distribution of the duration of whiffs and clean air and the distribution of the odorant concentration which were reported in the literature [2, 3]. Similarly to [32], we simulated plumes as pairs of odorant concentration time series, with a varying degree of correlation to emulate plumes of odors emitted from a single or from two separate sources (see [15] and Fig.S8). Similar to the previous section, the stimuli were applied to the models and we analyzed the PN responses in order to understand the ability of the early olfactory system to encode the signal. PN responses are very complex time series (see Fig.8) and many different decoding algorithms [42–46], could be present in higher brain areas to interpret them. However, as before, we applied the simple measure of peak PN activity in terms of the total firing rate above a given threshold to analyze the quality of the encoding.

**Fig 8.**
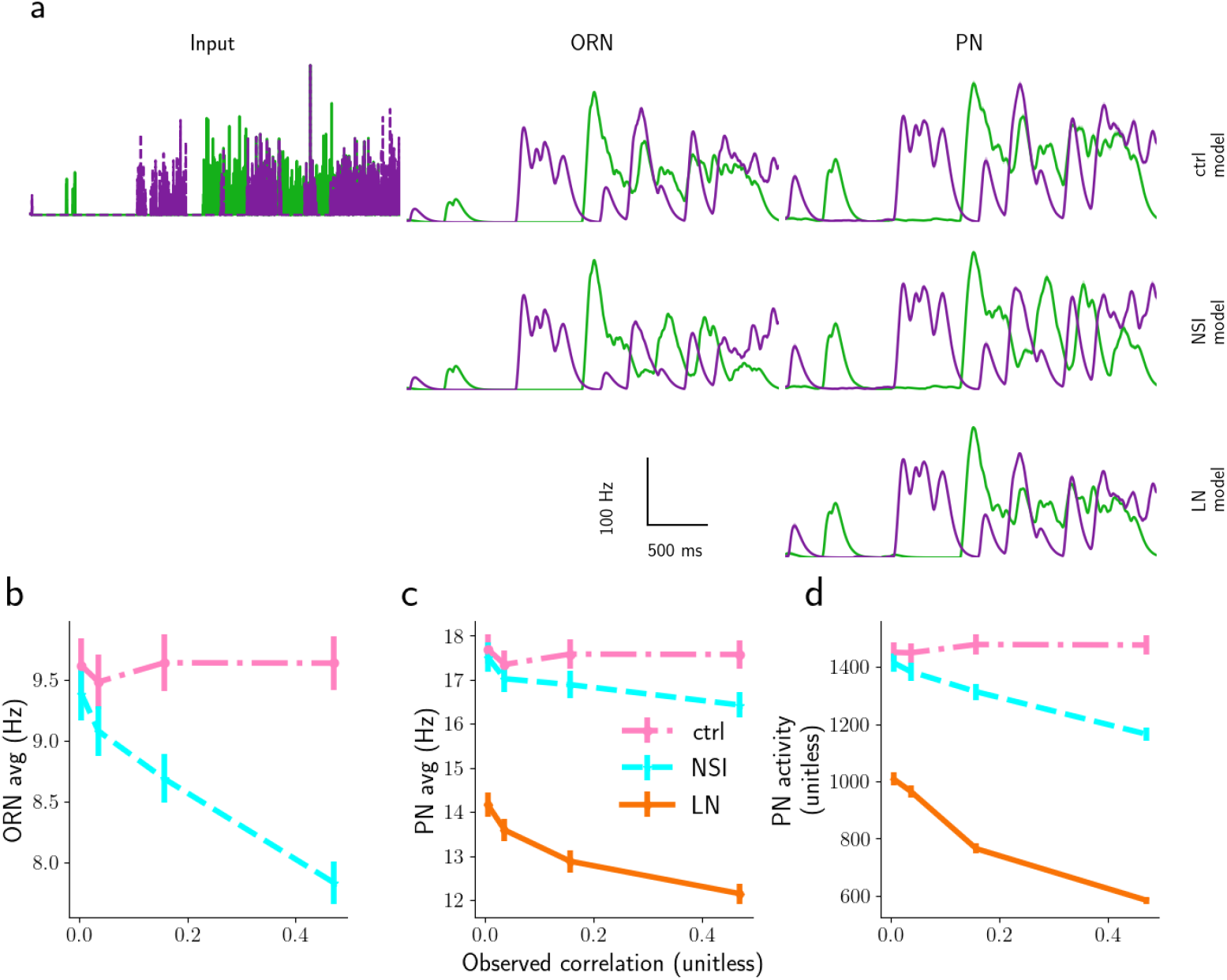
Responses to realistic plumes of the control, LN and NSI model. a) Time course of stimulus concentration (Input ORN, first column), and response of ORNs (second column), PNs (third column) to two 4 s long realistic plumes with statistical properties as described in the text; first row: control model, second row: NSI model, third row: LN model. Lines are the mean and shaded areas around the lines the standard deviation over 10 trials. b) Response of ORNs, and c) response of PNs averaged over 200 s for the three models: control model (dot dashed pink), LN model (orange continuous), and NSI model (dashed cyan). The observation from the LN model is not shown in panel b as it overlaps with the dot dashed pink lines (ctrl-model). d) Total PN activity above 150 Hz, for 3 ms maximum whiff durations.

To analyze the discrimination of plumes with odorants coming from a single source – highly correlated stimuli – from separate sources – poorly correlated stimuli – we developed a method to generate plumes of a prescribed correlation between concentration time series while keeping other properties such as intermittency and average odorant concentration constant (see Model and methods and SUPP Fig.S11). Using this method, we then explored plumes with correlation 0 to very close to 1 simulating the model for 200 s duration (a few times the maximal timescale in plumes, i.e. 50 s) and preset correlations between the odorant concentration time series, and first inspected the average activity of neuron types over the stimulation period. By construction, the ORN activity for the LN model is the same as in the absence of inhibition (8a-b), while the average ORN activity for the NSI model is lower and depends on the correlation between odor signals (8a-b). These effects are approximately the same for the whole range of the tested NSI strengths *ω*_*NSI*_. The situation is similar for the average PN activity. The average PN response in the NSI model is weakly dependent on input correlation (see 8c). The average PN responses in the LN inhibition network are lower than in the control model (8c). Hence, both mechanisms are useful for encoding input correlation with the average PN activity. All reported effects remain approximately the same for the entire range of explored parameters (*ω*_*NSI*_ and *α*_*LN*_).

Apart from the average activity, other measures can be used. A biologically plausible and commonly adopted hypothesis argues that information is conveyed in the high activity, i.e. firing rates above a given threshold. In this light we analysed the “peak PN” response, defined as the integrated PN activity over time windows where the PN firing rate is above a given threshold (e.g. 50, 100, or 150 ms). Fig. 8d shows peak PN for 150 Hz threshold (see SUPP Fig. S12 for plots with other underlying thresholds).

For the LN model and the NSI model, peak PN responses depend on the plume correlation. Within the values we investigated, the highest peak threshold of 150 Hz recovers the most information about input correlation and for high peak thresholds the NSI mechanism leads to more informative responses than LN inhibition. We conclude that most of the information about input correlations is contained in the first part of the response before adaptation takes place and that therefore the average activity over the entire response is not a good proxy for encoding the correlations in the input signals.

So far we have used simulated plumes corresponding to 60 m distance from the source. At different distances the maximum whiff durations will vary [29]. We therefore asked whether and how the efficiency of the two mechanisms depends on maximum whiff duration and hence distance from the source. To address this question, we generated plumes with different maximum whiff duration, *w*_*max*_. 9a shows a plot for each tested value of *w*_*max*_ (from 0.01 to 50 m) for peak threshold 150 Hz (see SUPP S14 and SUPP S15 for results with peak thresholds of 50, and 100 Hz). The choice of maximum whiff durations reflects typical experimental observations [2].

Two effects are evident: 1. At zero correlation between the stimuli, PN responses in the NSI model are quite similar to those in the control model while those in the LN model differ more, and 2. The PN responses in the NSI model depend more strongly on the input correlations of the stimuli than the PN activities in the LN model, especially for longer (>3s) whiffs (9a). This second effect is important because ideally we would like the PN responses to differ maximally between highly correlated plumes and independent plumes in order to discern the two conditions.

To quantify these effects we measured the following distances: 1. The distance between peak PN of the NSI model (or LN model) and of the control model at zero correlation, defined as 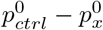 with *x* ∈ (NSI,LN) (9b) and 2. the distance between peak PN of NSI model (or LN model) at 0 correlation and at correlation (very close to) 1, defined as 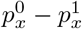 with *x* ∈ (NSI,LN) (9c). These figures show another clear advantage of using an NSI mechanism instead of LN inhibition for encoding correlation between odor plumes: 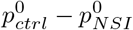 is always always smaller than 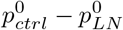 and 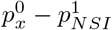 is consistently larger than 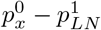.

However, the same measures for lower thresholds (i.e. 50 and 100 Hz) show that the two models are equivalent (see SUPP Fig.S14 and Fig.S15).

### 2.5 Increasing ORN Dynamic Range and sensitivity

Until now, we analysed the effects of the NSI on two neurons for two odorants for which they were independently sensitive to. We now move forward in generalizing one of the aspect that is taking into account the possibility that a single odorant can generate a response to both ORNs. In a theoretical study [14], Vermeulen and Rospars proposed that, in this situation, NSIs may be useful to alter the ORN’s dose-response. We tested this hypothesis by measuring the lower odor detection threshold and the total dynamic range of a pair of ORNs co-housed in a single sensillum and stimulated by a single odorant.

We can see in Fig.10a that for the NSI model, the dose-response curve of the more sensitive ORN presents a peak before plateauing at higher concentration. As already indicated by Vermeulen et al., this peak would prevent the correct encoding of the odorant intensity with the ORNs’ response. However, it could be a very fast layer whose unique aim is to detect the odorant at a specific intensity (for example, in our case between 1e-6 and 1e-4).

**Fig 9.**
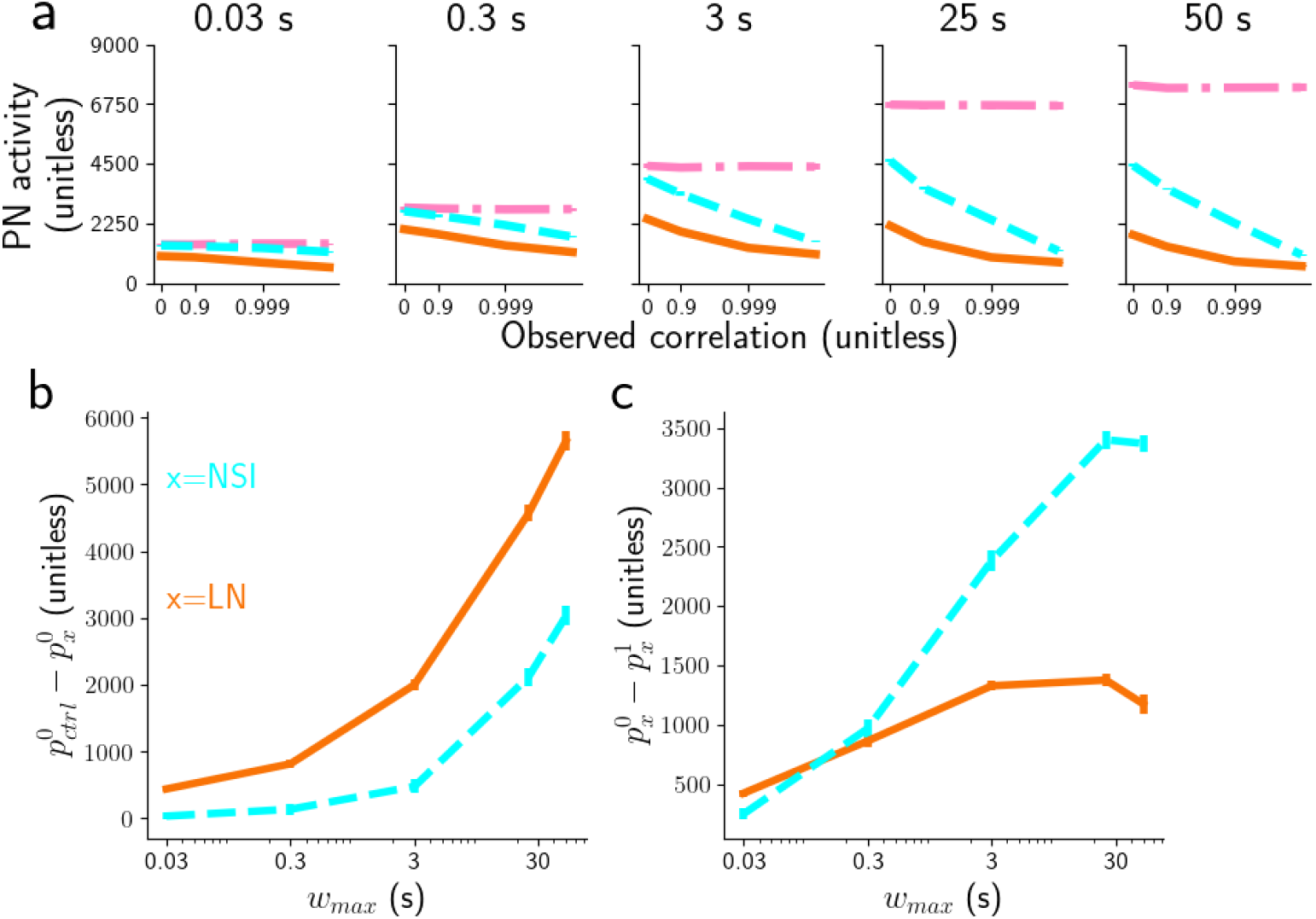
Comparing encoding efficiency of odorants’ correlation. a) Peak PN for threshold 150 Hz, and for different subsets of whiff durations (from 0.01 to 50s) for the three models: control model (dot dashed pink), LN model (orange continuous), and NSI model (dashed cyan). Note that the horizontal axis has a log-scale. b) Distance between the PN activity of the control model and the NSI model (or LN model), at 0 correlation, 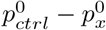 with *x* ∈ (NSI,LN). c) Distance between the PN activity of NSI model (or LN model) at 0 correlation and at correlation 1, 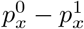 with *x* ∈ (NSI,LN).

**Fig 10.**
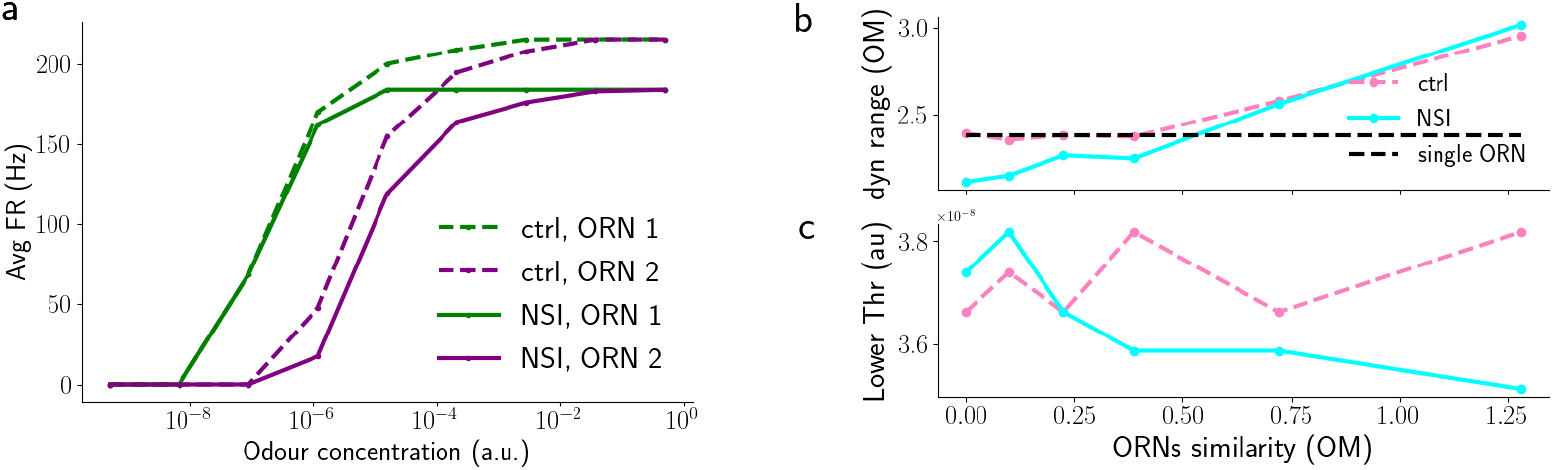
a) Dose response curves of two neurons for the control network (dashed lines) and for the NSI network (solid lines). b) Dynamic range of the two networks calculated for several values of the ratio in sensitivity of the two ORNs.

As for the first hypothesis tested, we used as stimuli short triangular pulses (10, 20, 50, 100 and 200 ms); the lower threshold is measured as the concentration value for which the ORN response reach 10% of its maximum (*C*_10%_) and dynamic range is measured as the commonly adopted definition of the logarithmic distance between the thresholds to 10% and to 90% of the sum of the dose-response curves of the two ORNs (i.e. log_10_(*C*_90%_*/C*_10%_), where *C*_*x*%_ is the concentration where the response reaches *x*% of the maximum response, see Fig. 10a). We measured them for pairs of ORNs that are increasingly similar. The ORNs similarity is measured as the logarithmic distance between the thresholds to 10% of the two isolated ORNs(i.e.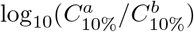, where a and b refer to the two ORNs). For this analysis, the comparison was between two ORNs interacting via NSI and two isolated ORNs (control).

Although our work differs in many aspects to the earlier theory (full temporal description of stimuli and the ORN and NSI implementation) and furthermore we took into consideration the total dynamic range and not simply of one ORN, our results are in line with the results reported by Vermeulen et al.: First, the NSI mechanism decreases the lower threshold for ORNs that respond to the same odors; Second, the NSI mechanism decreases the dynamic range for ORNs that respond to the same odors; Third, both lower threshold and dynamic range are best when responses of paired ORNs are completely different, or, indeed, NSIs are absent (see Fig.10b). Note that our control model is equivalent to the case where one ORN is not sensitive at all for the odorant in [14].

## 3 Discussion

*“Thought experiment is in any case a necessary precondition for physical experiment. Every experimenter and inventor must have the planned arrangement in his head before translating it into fact*.*”* E. Mach (1905)

We have implemented a model of the early olfactory system comprising ORNs of two receptor types, their NSIs in the sensillum, and two corresponding glomeruli in the AL, containing PNs and LNs in roughly the numbers that have been observed for *Drosophila*. Our objective was to investigate three potential roles of NSIs in insects’ olfactory processing: 1. Concentration invariant mixture ratio recognition, vital for insects to identify the type or state of an odor source (see e.g. [47–51] and references therein); 2. odor source separation, which can be critical for insects, e.g. in the context of finding mates [17, 18, 23]; 3. odor intensity perception [14].

By comparing our model with NSIs to a control model without lateral interactions between pathways for different receptor types, we found evidence that NSIs should be beneficial for concentration invariant mixture processing: NSIs lead to more faithful representation of odor mixtures by PNs in the sense that the ratio of PN activity is closer to the ratio of input concentrations when NSIs are present. Similarly, the mutual information between the ratio of input concentrations and the ratio of PN activity is higher for the NSI model. While we admittedly do not know how exactly odor information is represented in PN activity, responses that differ systematically with input ratio must be superior to responses that saturate and hence do not inform about the input ratio, as seen in the control model.

Furthermore, using a model variant with no NSIs but LN inhibition between glomeruli in the AL we found that 1. For synchronous individual whiffs, both models, the one with NSI mechanism and the one with LN inhibition, are better than the control model in several conditions (Fig.6 g); moreover, the NSI mechanism is typically more effective than LN inhibition. 2. For asynchronous individual whiffs, PN responses to the later whiffs are altered (inhibited) by the response to the first whiff when LN inhibition is present. This effect depends on the time scales of the LNs and the inhibitory synapses. In contrast, with NSIs the PN responses to the second whiffs are only mildly affected (increased) by the activity triggered by the first whiff. This effect is only evident for very high concentration, indicating that with NSIs there is less of a trade-off between benefits in encoding synchronous mixtures and distortions when odorants from separate sources mix.

These results further support the hypothesis that the NSI mechanism offers an evolutionary advantage by enabling more precise odor coding for these simple stimuli. Similar conclusions can be drawn when analyzing the capacity of the insect olfactory systems to encode the correlation between two odorants in a more realistic setting of an odor plume. We found that, when analysing peak PN activity (the integrated PN firing rate over windows during which it is above a given threshold), the model with NSI mechanism outperforms the LN inhibition model and both are better when considering peak activity than when considering average PN activity. Besides supporting the benefits of NSIs this also adds further evidence in favor of using peak activity to encode important features of a signal, in this case stimulus correlations, as hypothesized in earlier work (see e.g. [19, 52]).

The third hypothesis tested regards the idea illustrated in Fig.1b that the ORNs dynamic range could improve thanks to the NSIs [14, 25, 26]. The improvement of dynamic range by NSIs sits alongside work that showed that syntopic interactions at the receptor level and masking interactions at a cellular level achieve similar effects [53, 54] as well as improving mixture representations. Similarly, [55] showed that syntopic interactions improve concentration invariant mixture representation in particular for odors with many components. How these receptor-level and cell-level mechanisms interact with sensillum-level NSIs is an interesting future research question.

However, our results indicate the opposite: The total dynamic range of two isolated ORNs is greater than that of two ORNs interacting through NSIs. The disruptive effect of NSIs arises from the dependence of the NSI strength on the activity of the more strongly activated ORN, so that instead of preventing the ceiling effect for high concentrations, NSIs lead to an activity peak for central values of the concentration. The effect increases with the similarity between the sensitivity of the ORNs and in the limit of identical ORNs it is so strong that the resulting dynamic range is inferior to the dynamic range of a single ORN.

Furthermore, the non-monotonic peak-and-plateau response of the more strongly activated ORN degrades intensity encoding. However, the pair of neurons would be able to rapidly detect odorants at specific intensities.

### 3.1 The model and its limitations

*“A good model should not copy reality, it should help to explain it”*, [56].

As in every modelling work the level of description must match the purpose of the investigation. In terms of Marr’s categorisation of models [57], our model is somewhere between the algorithmic level - as both our models implement a form of lateral inhibition - and the implementation level - albeit we are not yet able to capture the underlying physics of the NSI mechanism. Because of our hypothesis that the role of NSIs is to improve processing of temporally complex stimuli, we focused on a description which included temporal dynamics but was otherwise as simple as possible. We therefore have simplified 1. the cellular dynamics of odor transduction [31, 58–62] and only heuristically describe the macroscopic effects at the receptor neuron level, an approach similar to [30]; 2. the complexity of the full receptor repertoire in the insect olfactory system, e.g. about 60 ORN types in *Drosophila*, and instead focused on a single sensillum with two co-housed ORNs; 3. the true complexity of the many different LN types and transmitters in the AL [63], using only GABA_A_-like LNs. 4. the spatial distribution of the sensilla on the surface of the antenna or the maxillary palp; 5. the complexity of odor stimuli delivered by stimulation devices in the experiments we are mimicking for the single pulse investigation (see the corresponding Model and methods section, [29]), 6. the asymmetry of NSIs where there is some evidence that the strength of the NSIs is proportional to the size of the ORN that is exerting the interaction onto another neuron [13]. By making these simplifications we were able to reduce the number of free parameters in the model, reasonably constrain most parameters and scan the few remaining parameters, such as the strength of LN inhibition, across a reasonable range. This increases our confidence that the observed benefits of NSIs for olfactory information processing are not artefacts of particular parameter choices in the model(s).

### 3.2 The future: how to generalize the model

*“[*…*] I have the solution, but it works only in the case of spherical cows in a vacuum*.*”*,

For the sake of simplicity we chose to work with a specific animal model in mind and because of the large amount of information available in the literature, we chose *Drosophila*. It will be interesting to see whether and how much our results can be generalized to other insects such as other flies or moths (that have 1-4 ORNs per sensillum), or even bees, ants and beetles (with up to 20-30 co-housed ORNs).

We can only speculate that all the effects of the NSIs shown here will be, at least partially, amplified, as there will be a much higher probability that the ORNs would be sensitive to the odorants present in odor mixtures. But of course, we have to consider the disruption generated by the decreased dynamic range whenever co-housed ORNs respond to the same odorants.

However, the complexity of this problem is clearly enough to require an extensive study that we hope can start from this work and the model that is freely available online.

### 3.3 Comparison with related modelling works

*“If I have seen further it is by standing on the shoulders of Giants*.*”* I. Newton (1675).

Our work builds on ideas in previous models (e.g., [14, 55, 64]) and concurrent approaches (e.g. [30]). While earlier modeling works focused on the oscillatory and patterned dynamics of activity in the antennal lobe [65–70], it was soon realized that the recognition of odorants and their mixtures across different concentrations posed a particularly difficult question. One school of models explored the idea of winnerless competition as a dynamical systems paradigm for concentration invariant coding [71, 72] while others explored more direct gain control mechanisms mediated by local neurons in the AL [35, 36, 73]. The task becomes even more difficult when the exact ratio of mixtures needs to be recognised, and a network model for mixture ratio detection for very selective pheromone receptors has been formulated in [37]. However, generally, odors already interact at the level of individual ORs due to competitive and non-competitive mechanisms which can be recapitulated in models, see e.g. [64] for vertebrates and [55] for invertebrates.

However, our model also makes a clear departure from the large number of models that have been built on assumptions and data based on long, essentially constant, odor step stimuli. While these kind of stimuli are not impossible, they can be considered as the exception more than the rule; for instance, even at more than 60 m from the source, around 90% of whiffs last less than 200 ms [1, 74], see [29] for review. This insight is particularly difficult to reconcile with models that emphasize and depend on intrinsically generated oscillations in the antennal lobe [65–70], and models that depend on comparatively slow, intrinsically generated dynamics such as the models based on the winnerless competition mechanism [71, 75]. The original interpretation of these models, how they use intrinsic neural dynamics to process essentially constant stimuli, is disrupted when stimuli have their own fast dynamics. How to reconcile the idea of intrinsic neural dynamics for information processing with natural odor stimuli that have very rich temporal dynamics of their own remains an open problem.

In building our model, we followed the main ideas developed by [14] but went beyond the assumption of constant stimuli and also added the important element of adaptation in ORNs and PNs, a widely accepted feature that is important in the context of dynamic stimuli. While Vermeulen et al. were already interested in possible evolutionary advantages of NSIs (without finding much), we here added the comparison with lateral inhibition in the AL that has been described as a competing mechanism, from an experimental (e.g. [11]) and a theoretical point of view (e.g., [35–37, 76]. Finally an important addition in this study are the mixture stimuli: Many, though not all, earlier works focused on the response of the network to mono-molecular odors, whereas we analyse the network response to two-odorant mixtures.

A previous study with very similar motivation relating to mixture ratio recognition is the analysis of pheromone ratio recognition of [37]. However, this earlier work still assumed constant stimuli, no adaptation in ORNs, a fixed target input ratio and only LN inhibition.

### 3.4 Further hypotheses about NSIs

*“There is always a well-known solution to every human problem—neat, plausible, and wrong*.*”* H. L. Mencken 1920 “Prejudices: Second Series”

At this early stage, our knowledge and understanding of NSIs is still full of gaps. For example, while suggestive our results cannot prove beyond doubt whether NSIs are effectively useful to the olfactory system, or whether they are just an evolutionary spandrel. We also do not know their evolutionary history. One interesting idea would be that the complex function of improved odor mixture encoding could have arisen as a side effect from a simpler function, e.g. of saving space, but we do not have any evidence to support this.

Researchers in the past 20 years have suggested a number of non exclusive explanations for the functions of NSIs. We have analyzed three of them - improved odor ratio representation, and improved dynamic range, and detecting plume correlations. An alternative hypothesis is that NSIs could facilitate novelty detection for odor signals on the background of other odors [11], if newly arriving “foreground odors” suppress the ongoing response to an already present “background odor”. Todd et al. already noticed that NSIs duplicate the role of LNs in the AL even though [7] pointed out later that LN networks take effect later and mainly decorrelate PN activities and normalize them with respect to the *average input* from ORNs. Here we have added to the discussion by showing that NSIs have advantages with respect to their faster timescale that leads to less disruption of asynchronous odor whiffs.

Moreover, NSIs have two additional key advantages with respect to LN inhibition in the AL or processes in later brain areas: 1. NSIs can be energetically advantageous as they don’t need to generate spikes [6, 77–81]. 2. NSIs take place at the level of the single sensillum and hence a few spikes and synapses earlier than any AL or later interactions [7, 11]. In the AL the information from ORNs of the same type is likely pooled and information about the activity of individual ORNs is not retained (see e.g. [40, 82]). Therefore, while interactions within the sensillum are precise in space and time, interactions in the AL will be global (averaged over input from many sensilla) and information channels will interact in an averaged fashion. Similar local interactions in the very early stages of sensory perception were already discussed for the retina [83, 84].

### 3.5 Conclusions

In conclusion, we have demonstrated in a model of the early olfactory system that NSIs have advantages over LN inhibition in the AL with respect to faithful mixture ratio recognition and plume separation. In our future work we seek to confirm the behavioral relevance of NSIs in *Drosophila*. Other interesting future directions include the relationship of NSIs and syntopic effects/masking, as well as the differential roles of NSIs and LN inhibition when both are present at the same time.

## 4 Model and methods

### 4.1 Model topology

We model the electrical activity of the early olfactory system of *Drosophila melanogaster*. The model encompasses ORNs on the antenna, and the matching glomeruli in the AL, containing PNs and LNs. For simplicity, ORNs are housed in sensilla in pairs, and each neuron in a pair expresses a different OR type. The paired neurons interact through NSIs, effectively leading to mutual inhibition (see Fig.1 a). There are multiple sensilla of the same type on each antenna. We here model 40 sensilla per type [82]. ORNs of the same type all project exclusively to the same glomerulus in the AL, making excitatory synapses onto the associated PNs. In addition to the inputs from ORNs, PNs also receive global excitation from PNs associated with other glomeruli and from other parts of the brain. They are inhibited by the LNs of other glomeruli but not by LN in the same glomerulus (see Fig.2). The model simulates one type of sensillum and hence two types of ORNs, ORN_*a*_ and ORN_*b*_. We assume that ORN_*a*_ and ORN_*b*_ are selectively activated by odorants A and B, respectively (see Fig.2 and Fig.1 a).

### 4.2 Olfactory Receptor Neurons

We describe ORN activity in terms of an odorant transduction process combined with a biophysical spike generator [30]. During transduction, odorants bind and unbind at olfactory receptors according to simple rate equations. As we are not interested in the competition of different odorants for the same receptors, we simplify the customary two-stages binding and activation model [55, 64, 85] to a single binding rate equation for the fraction r of receptors bound to an odorant,

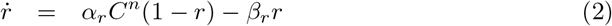

We described the spike generator by a leaky integrate-and-fire neuron with adaptation,

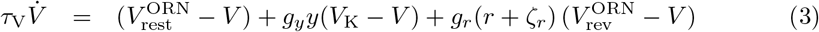

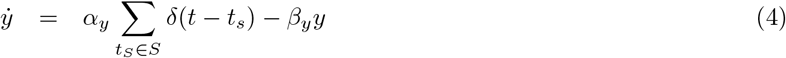

where, *V*_rest_ is the resting potential, *V*_rev_ the reversal potential of the ion channels opened due to receptor activation, *ζ*_*r*_ is Gaussian colored noise with zero mean and standard deviation *r*_noise_, representing receptor noise, and *y* is a spike rate adaptation variable with decay time constant *b*_*y*_.

The parameters *a*_*y*_ and *b*_*y*_ are estimated together with *b*_*r*_ and *d*_*r*_ to reproduce the data presented in [28]. The model is similar in nature to the models presented in [30, 86] albeit simplified and formulated in more tangible rate equations. As we will demonstrate below, this simplified model can reproduce experimental data equally well as the previous models.

### 4.3 Non-synaptic interactions

To simulate the NSI, we assumed that the electric interaction takes place through the reversal threshold and that it would change linearly with the activation variable of the co-housed neuron. Omitting ORN in the subscript, we obtain the following equations:

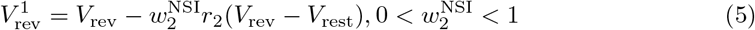

### 4.4 The antennal lobe

We here reduce the antennal lobe (AL) to two glomeruli, a and b (see Fig.2) in order to focus on the effects of NSIs of the corresponding ORN types. The numbers of PNs and LNs per glomerulus are in qualitative agreement with what reported in literature [82, 87–89].

The competing LNs are inhibitory whereas the PNs are excitatory. For simplicity, we do not model multiple kinds of LNs or PNs that have been observed in the AL. Similar models are being used extensively in the analysis of the insect AL [36, 37, 55, 73] and are well suited for replicating the competition dynamics that we seek to evaluate.

We model neurons as leaky integrate-and-fire (LIF) neurons with conductance based synapses. For PN *i*,

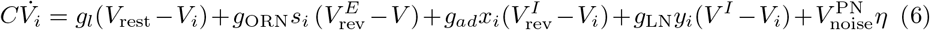

where *V*_*i*_ is the membrane potential of PN 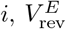 and 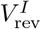 are reversal potential for the excitatory and the inhibitory input, respectively, *s*_*i*_ is the combined activation of synaptic inputs from connected ORNs, *x*_*i*_ is the activation of a spike rate adaptation current, *y*_*i*_ is the activation of inhibitory synapses from connected LNs and *η* Gaussian white noise with zero mean and standard deviation one. The LNs are described by a similar model but without adaptation,

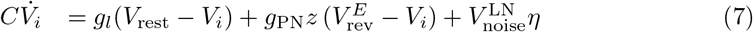

where *V*_*i*_ is the membrane potential of LN *i*, and *z* represents the activation of excitatory synapses from the PN. For both PNs and LNs, when V> Θ, the neuron fires a spike and V is reset to V_rest_ and it does not change for a refractory period, *τ*_ref_, lasting 2 ms. Synaptic activation is modeled according to

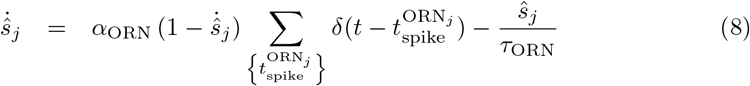

where the sum is over all spikes in ORN *j*, and the resulting overall activation *s*_*i*_ for PN *i* is 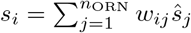 where *w*_*ij*_ is the connectivity matrix between ORNs and PNs, *w*_*ij*_ = 1 if ORN *j* is connected to PN *i* and *w*_*ij*_ = 0 otherwise. Similarly, for PN to LN excitation,

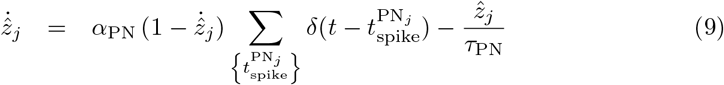

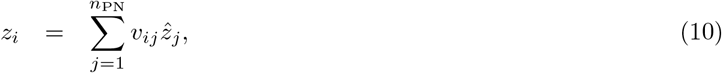

with connectivity *v*_*ij*_, for LN to PN inhibition,

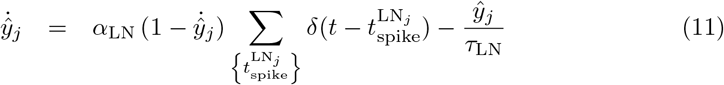

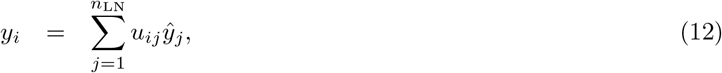

with connectivity *u*_*ij*_, and for the activation of the spike rate adaptation current,

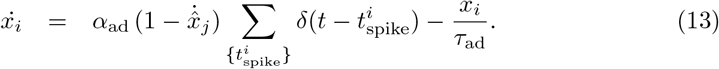

Each one of these variables has its time constant - *τ*_*s*_, *τ*_*V*_, *τ*_ad_, and *τ*_*y*_. The multiplicative factors *α*_*LN*_, *α*_*ORN*_ reflects the amount of released vesicles per each spike from an ORN and LN, respectively and they can be considered synaptic strength.

All the parameters used for the simulations are reported in Table 1. The comparative analysis between the LN inhibition and NSI mechanism has been carried out through the exploration of the two parameters *α*_*LN*_ and the strength of the NSIs, *ω*_*NSI*_.

The simulations were run with custom made Python code available online at *https://github.com/mariopan/flynose*.

## 5 SUPPLEMENTARY MATERIALS

**Fig S1.**
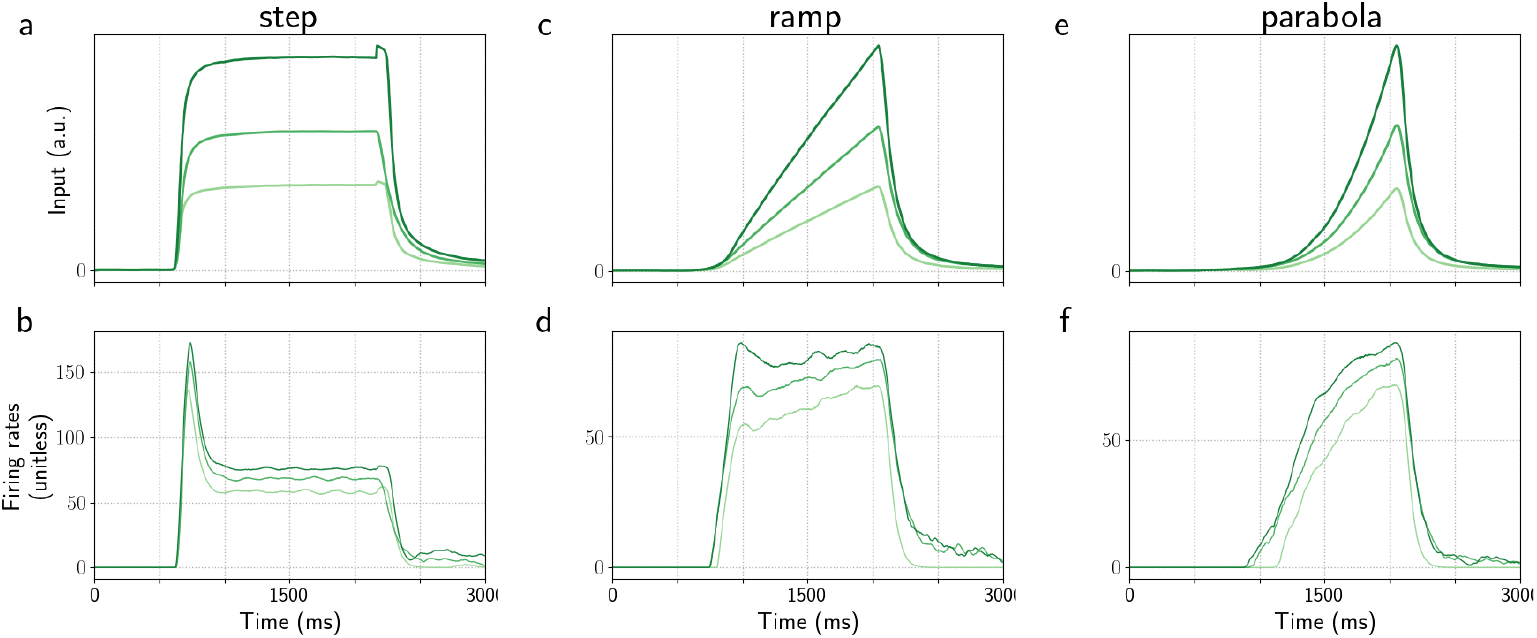
Model ORN response to a single step, a ramp, and a parabola as in [30]. Model ORN response to a single step (a,b), ramp (c,d), and parabola (e,f). a, c, e: Stimulus waveforms, i.e. odorant concentration profiles, as in [38]. b, d, f: Model ORN firing rates visualized as a spike density function (SDF).

**Fig S2.**
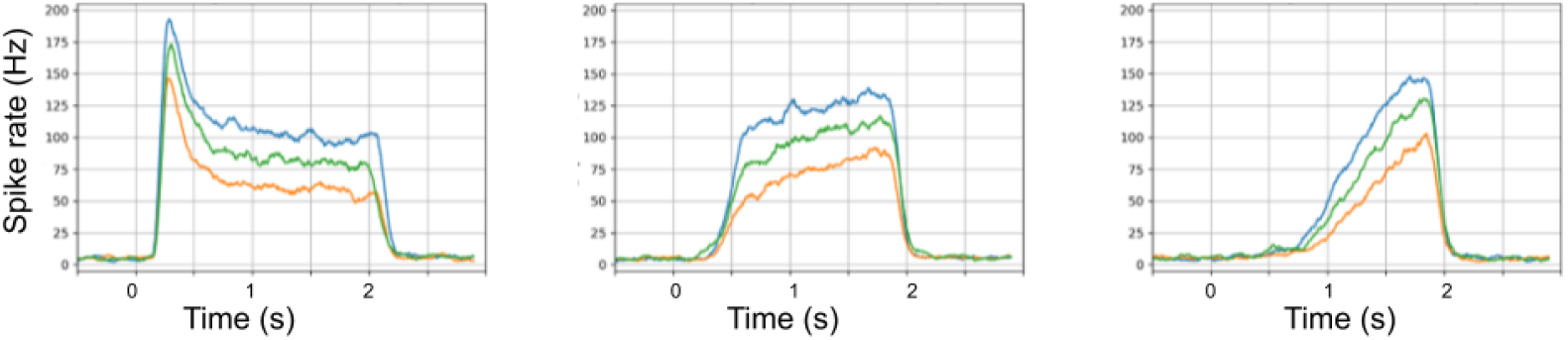
Output of the model of Lazar and Yeh [30] for the Or59b receptor neuron in response to the corresponding stimulus waveforms (experimental data reported in [38]).

**Fig S3.**
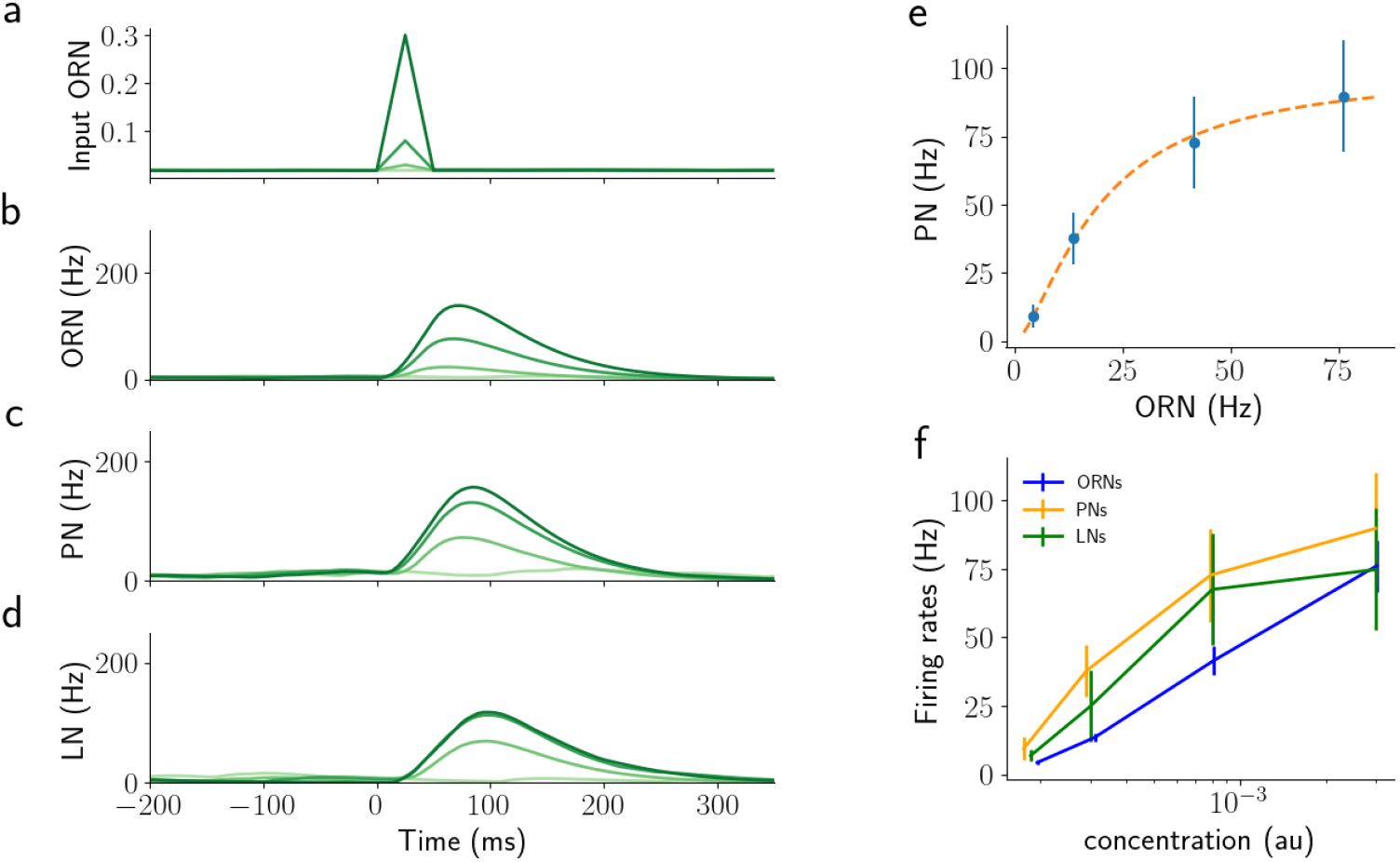
Similar results of Fig.4 for shorter stimulation time (50 ms). a) 50 ms step stimuli, shade of green indicates concentration. b)-d) corresponding activity of ORNs, PNs, and LNs. Shades of green match the input concentrations. e) Average response of PNs over 50 ms against the average activity of the corresponding ORNs. The orange dashed line is the fit of the simulated data using equation eq. 1 as reported in [41]. f) Average values for PNs, ORNs, and LNs for different values of concentration. Error bars show the SE over PNs.

**Fig S4.**
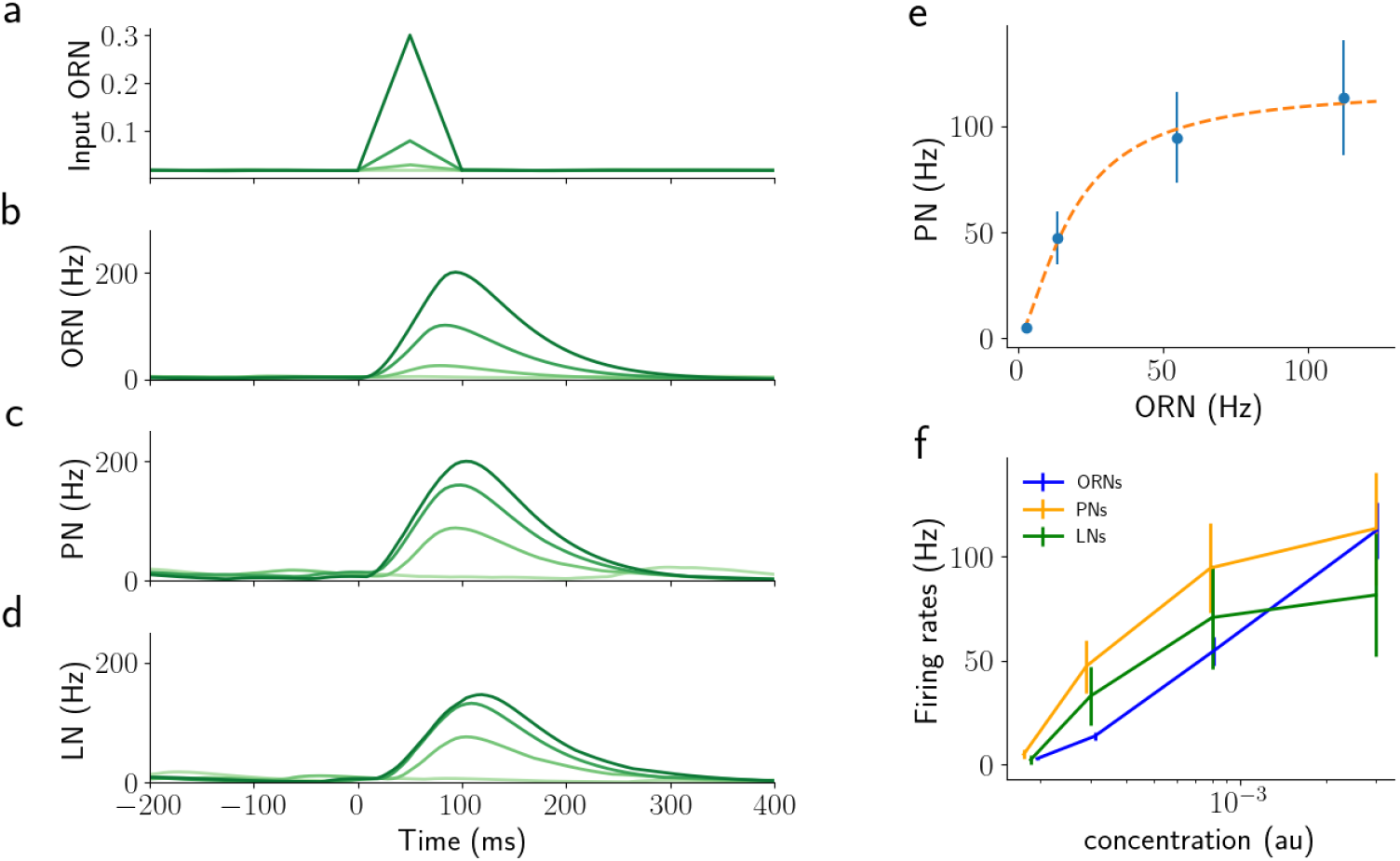
Similar results of Fig.4 for shorter stimulation time (100 ms). a) 100 ms step stimuli, shade of green indicates concentration. b)-d) corresponding activity of ORNs, PNs, and LNs. Shades of green match the input concentrations. e) Average response of PNs over 100 ms against the average activity of the corresponding ORNs. The orange dashed line is the fit of the simulated data using equation eq. 1 as reported in [41]. f) Average values for PNs, ORNs, and LNs for different values of concentration. Error bars show the SE over PNs.

**Fig S5.**
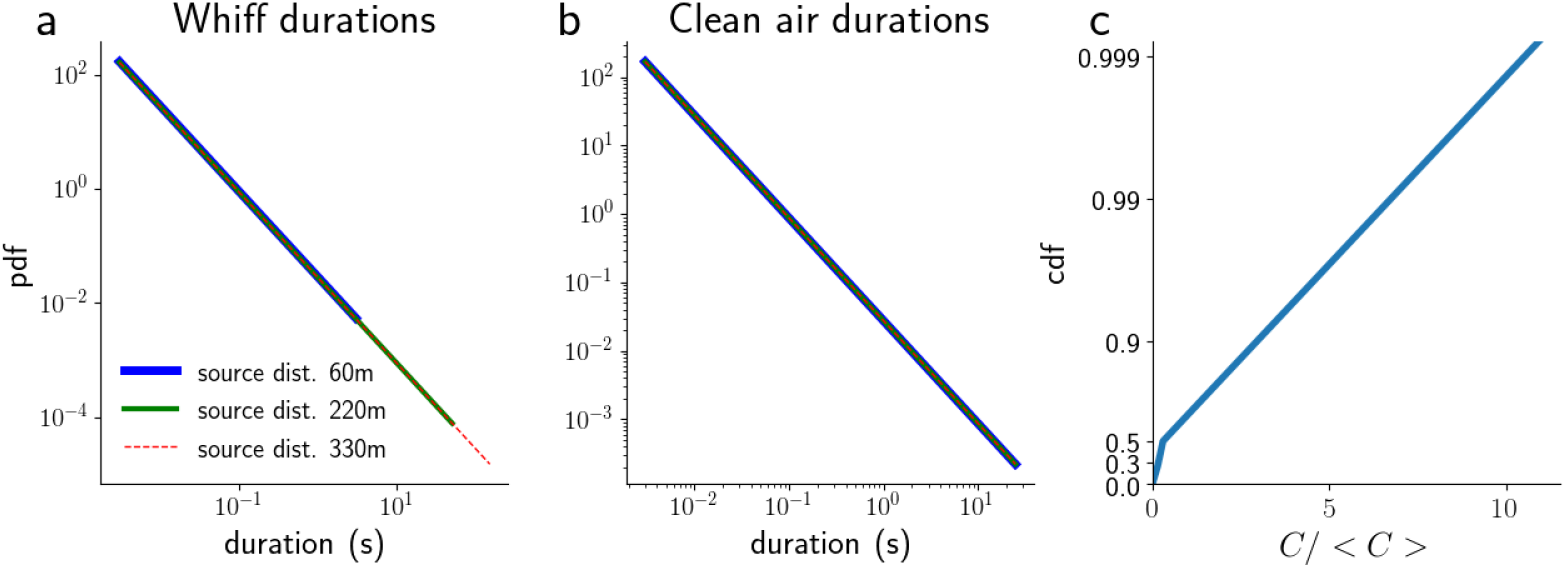
Plume statistics of natural plumes. a) Probability distribution of the whiff durations for odorants emitted at distances larger than 60 m [2]. b) Probability distribution of the blank durations for odorants emitted at distances larger than 60 m [2]. c) Probability distribution of the normalized concentration for odorants emitted at 75 m distance from the source [3].

**Fig S6.**
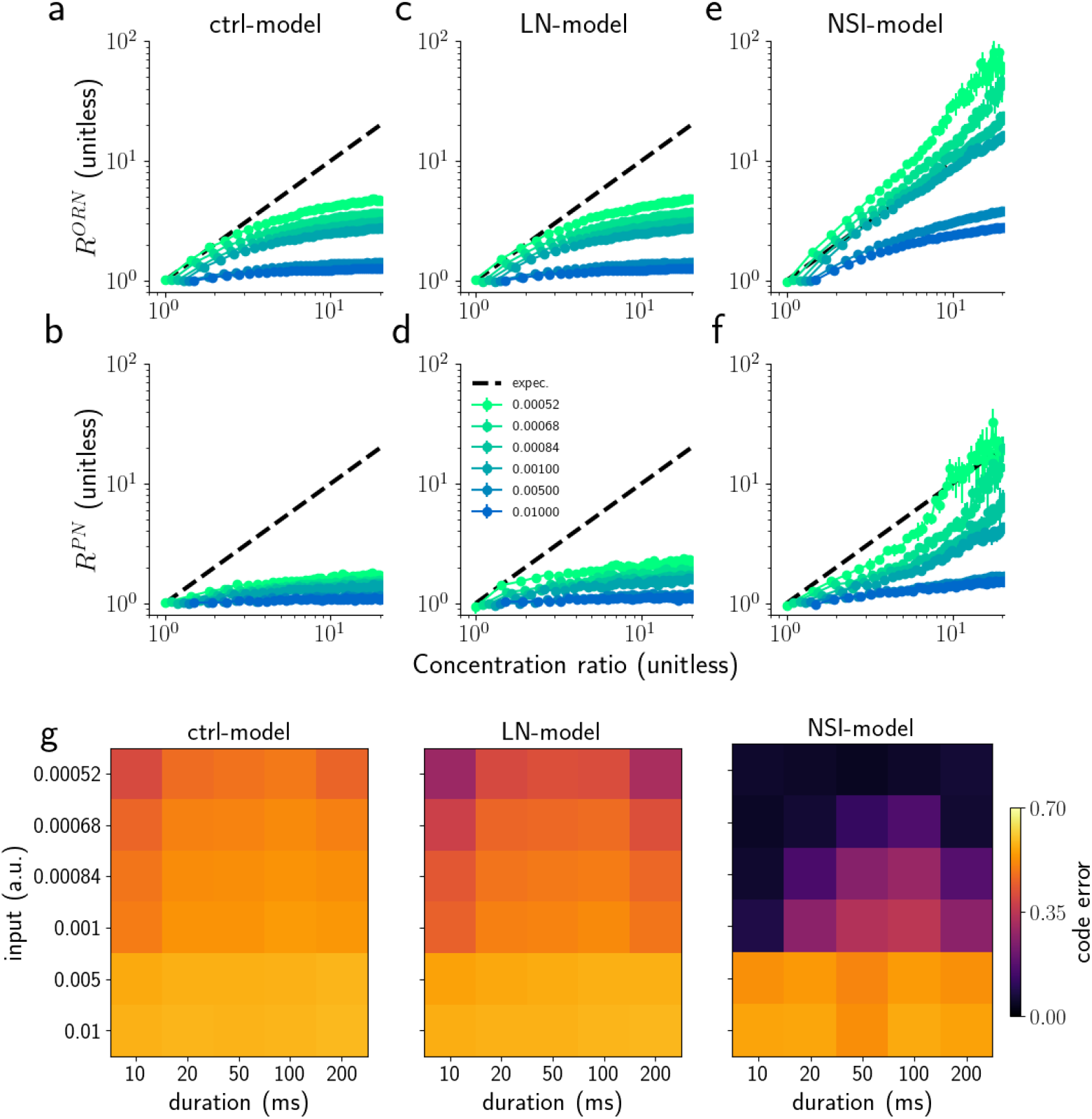
Encoding ratio with the average PN activity. Encoding ratio with the average PN activity. ORN (a,c,e) and PN (b,d,f) responses to a single synchronous triangular pulse of 50 ms duration applied to both ORN groups. The graphs show average responses ratio (*R*^*ORN*^ and *R*^*P N*^), respectively, versus concentration ratio of the two odorants for four different overall concentrations (colours, see legend in f). The average PN responses would be a perfect reflection of the odorant concentration if they followed the black dashed diagonal for all concentrations. Error bars represent the semi inter-quartile range calculated over 50 trials. g) Analysis of the coding error for different values of stimulus duration (from 10 to 200ms) and concentration values (0.2 to 1.4).

**Fig S7.**
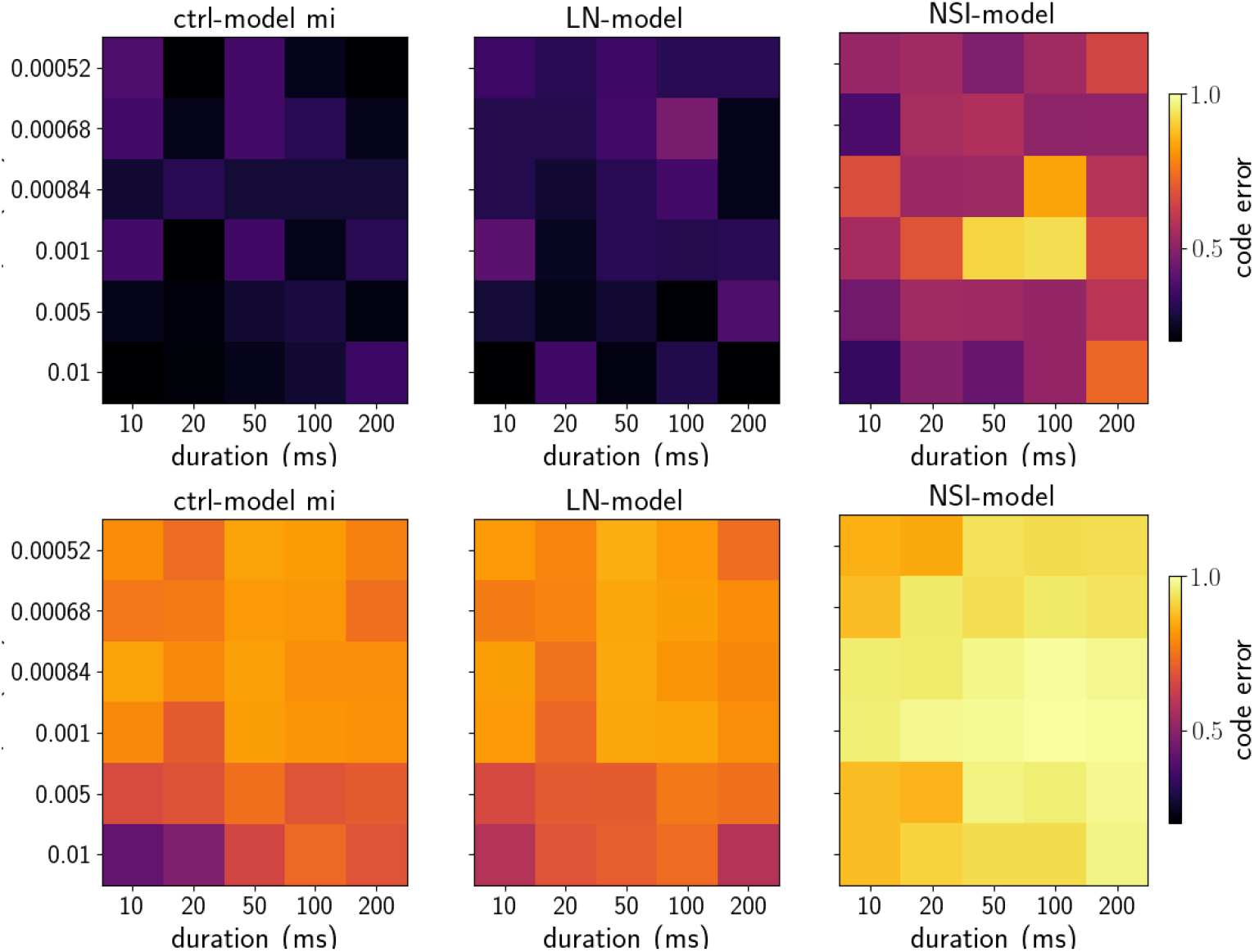
Encoding ratio analysis with MI. Analysis of the coding error with mutual information for different values of stimulus duration (from 10 to 200ms) and concentration values. ON TOP) The coding error is calculated as the MI between odorant concentration and *R*^*P N*^ (see Model and methods). ON THE BOTTOM) The coding error is calculated as the Pearson correlation between odorant concentration and *R*^*P N*^ (see Model and methods)..

**Fig S8.**
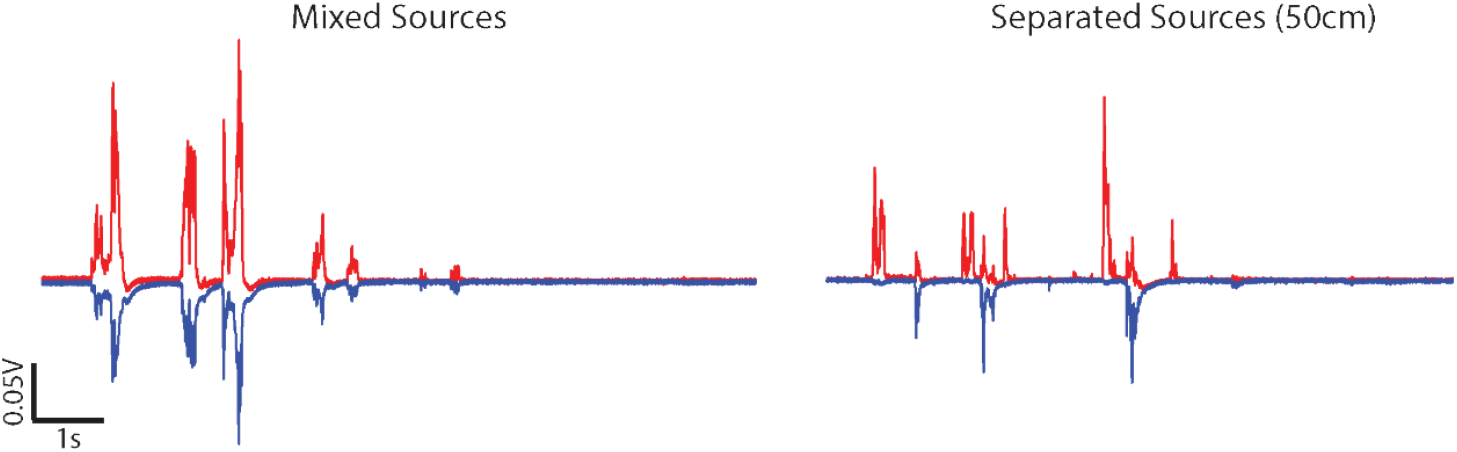
Example concentration fluctuation time series of natural plumes for two odorants emitted by a single source or two separate sources [15].

**Fig S9.**
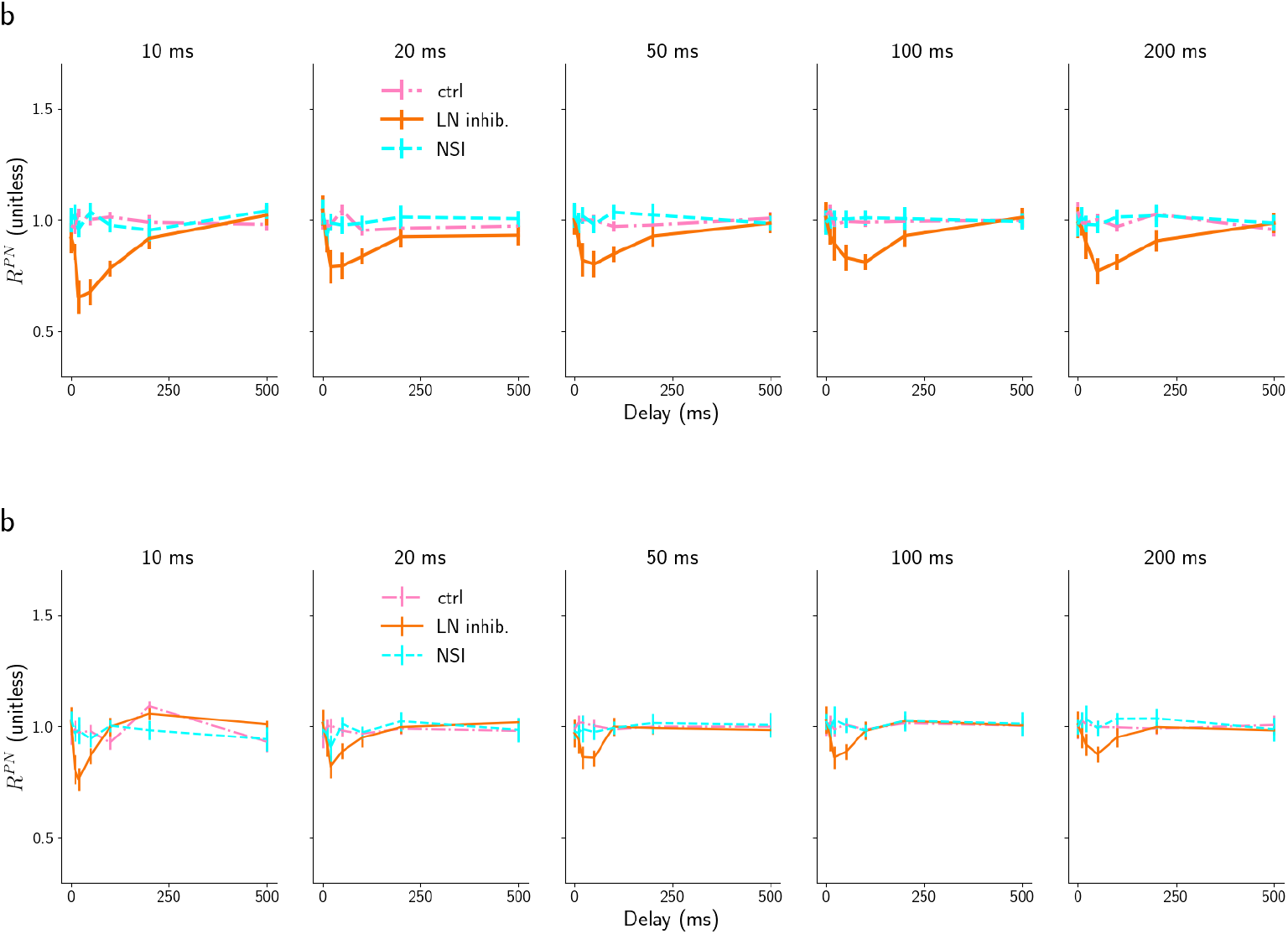
Results for the average PN response over the stimulus duration. Median ratio of the average PN responses of the two glomeruli 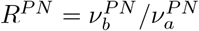 in the three models:control model (dot dashed pink), LN model (orange continuous), and NSI model(dashed cyan) for different stimulus durations as marked on the top. Panel a) shows the results using *τ*_*LN*_ = 250*ms* and panel b) *τ*_*LN*_ = 25*ms*. Error bars represent the semi inter-quartile ranges.

**Fig S10.**
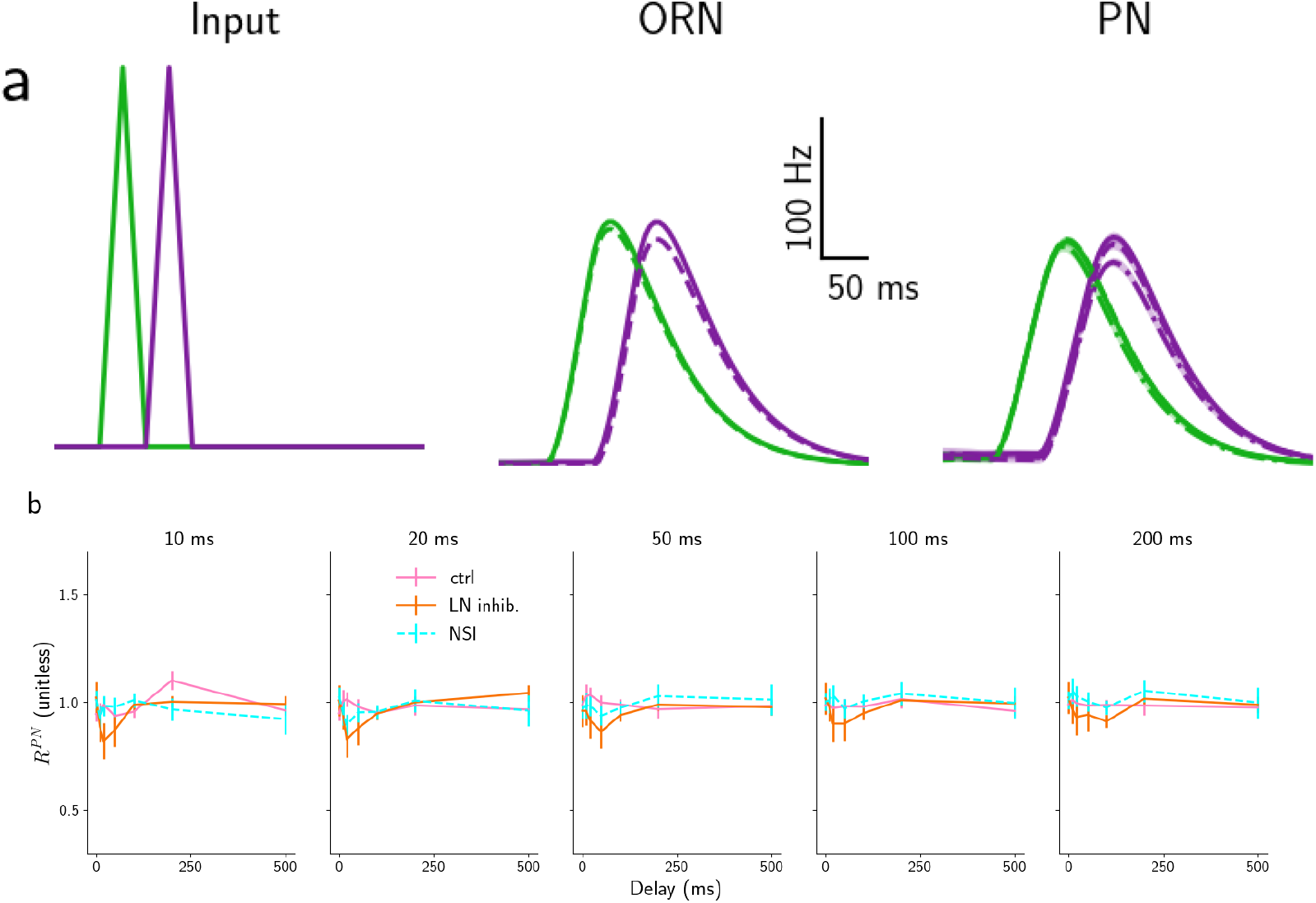
Responses to asynchronous pulses of the control, LN and NSI model. Panel a) Time course of ORN and PN activity in response to two asynchronous triangular pulses (50 ms, 1st column) for the three models – control (continuous line), NSI (dashed line) and LN model (dot and dashed line). Input to the three models is identical, while control and LN models have identical ORN activity, which is therefore not displayed twice. The two colours represent the two odors, ORN and PN types. The lines show the average response and the shaded area around the lines the standard deviation over 10 trials. Panel b) Median ratio of the peak PN responses of the two glomeruli 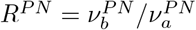 in the three models: control model (dot dashed pink), LN model (orange continuous), and NSI model (dashed cyan) for different stimulus durations as marked on the top. Error bars represent the semi inter-quartile ranges. The same plots are shown for *τ*_*LN*_ is low (250 ms) in Fig.7.

**Fig S11.**
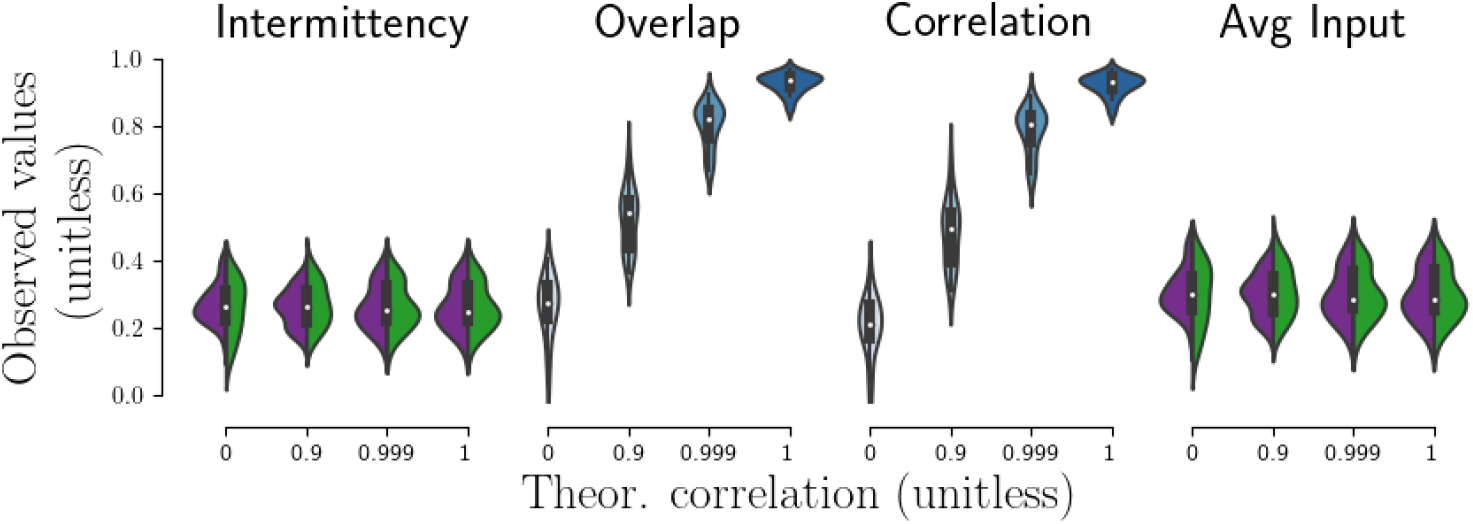
Statistical properties of simulated natural plumes. Observed properties of the simulated plumes as a function of the intended correlation between plumes averaged over 200 s. Intermittency and average input plots show the values for the two plumes (green and purple).

**Fig S12.**
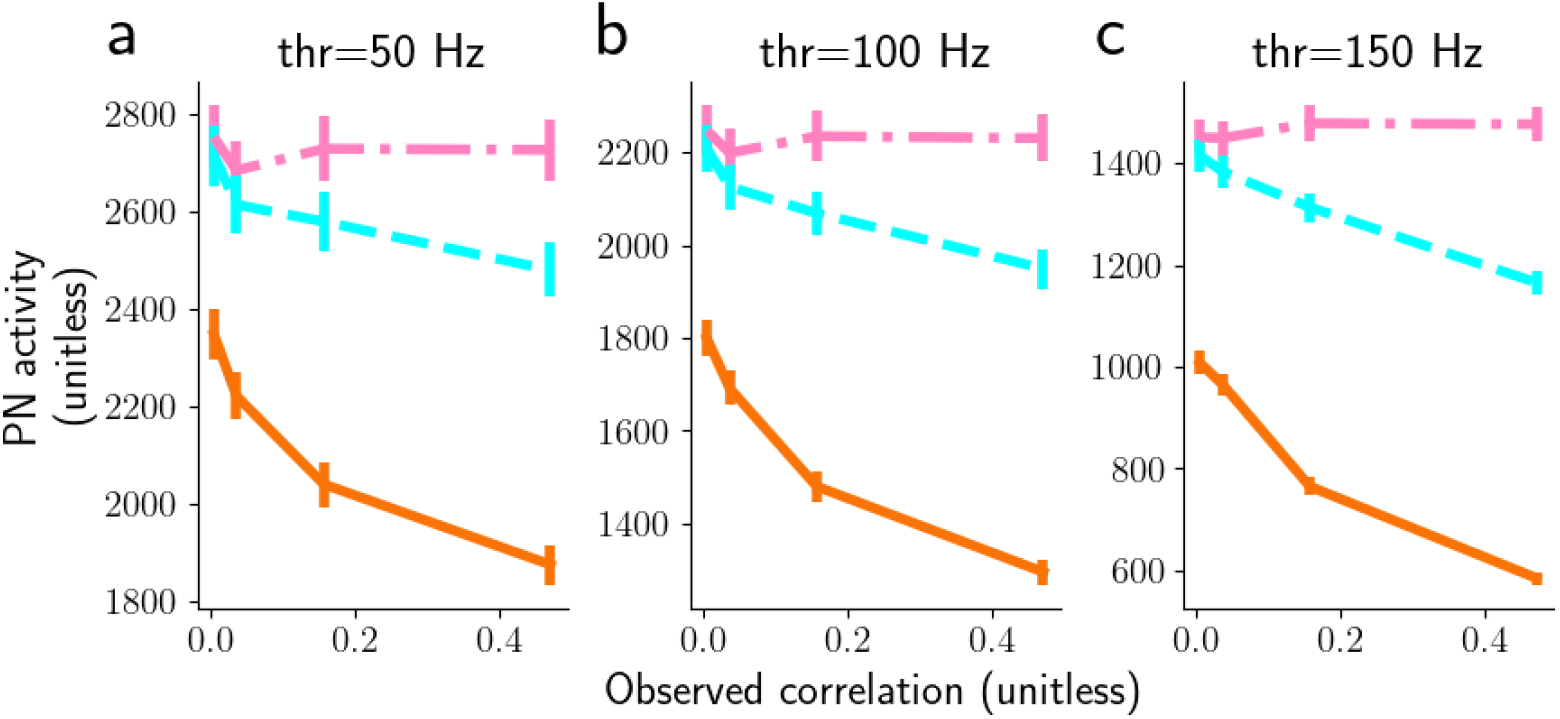
Similar results of Fig.8d with different thresholds (50, 100, 150 Hz). Panels a-c) show the total PN activity above 50, 100, 150 Hz, respectively, for 3 ms maximum whiff durations.

**Fig S13.**
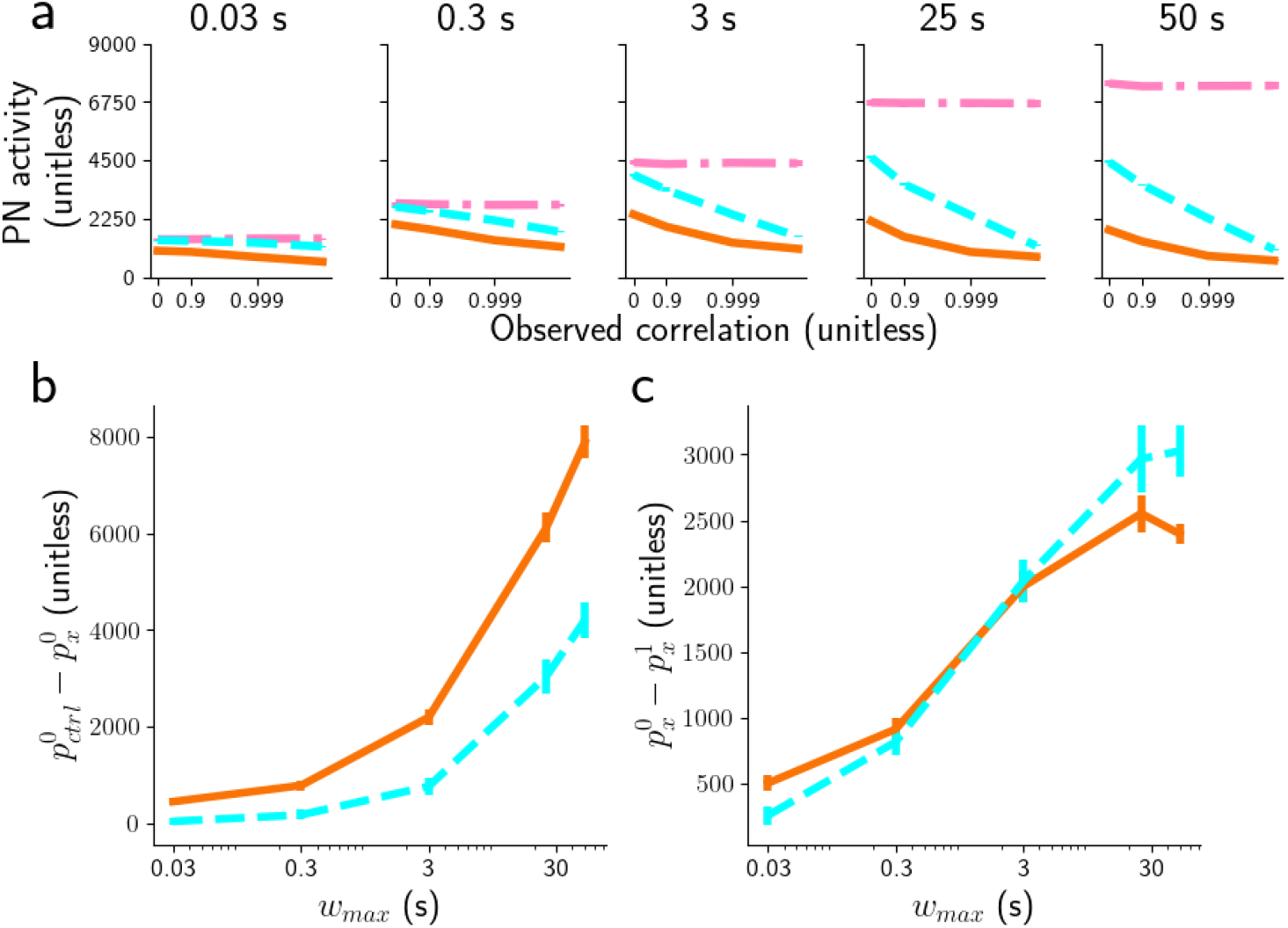
a) Peak PN for threshold 150 Hz, and for different subsets of whiff durations (from 0.01 to 50s) for the three models: control model (dot dashed pink), LN model (orange continuous), and NSI model (dashed cyan). Note that the horizontal axis has a log-scale. b) Distance between the PN activity of the control model and the NSI model (or LN model), at 0 correlation, 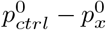 with *x* ∈ (NSI,LN). c) Distance between the PN activity of NSI model (or LN model) at 0 correlation and at correlation 1, 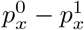 with *x* ∈ (NSI,LN).

**Fig S14.**
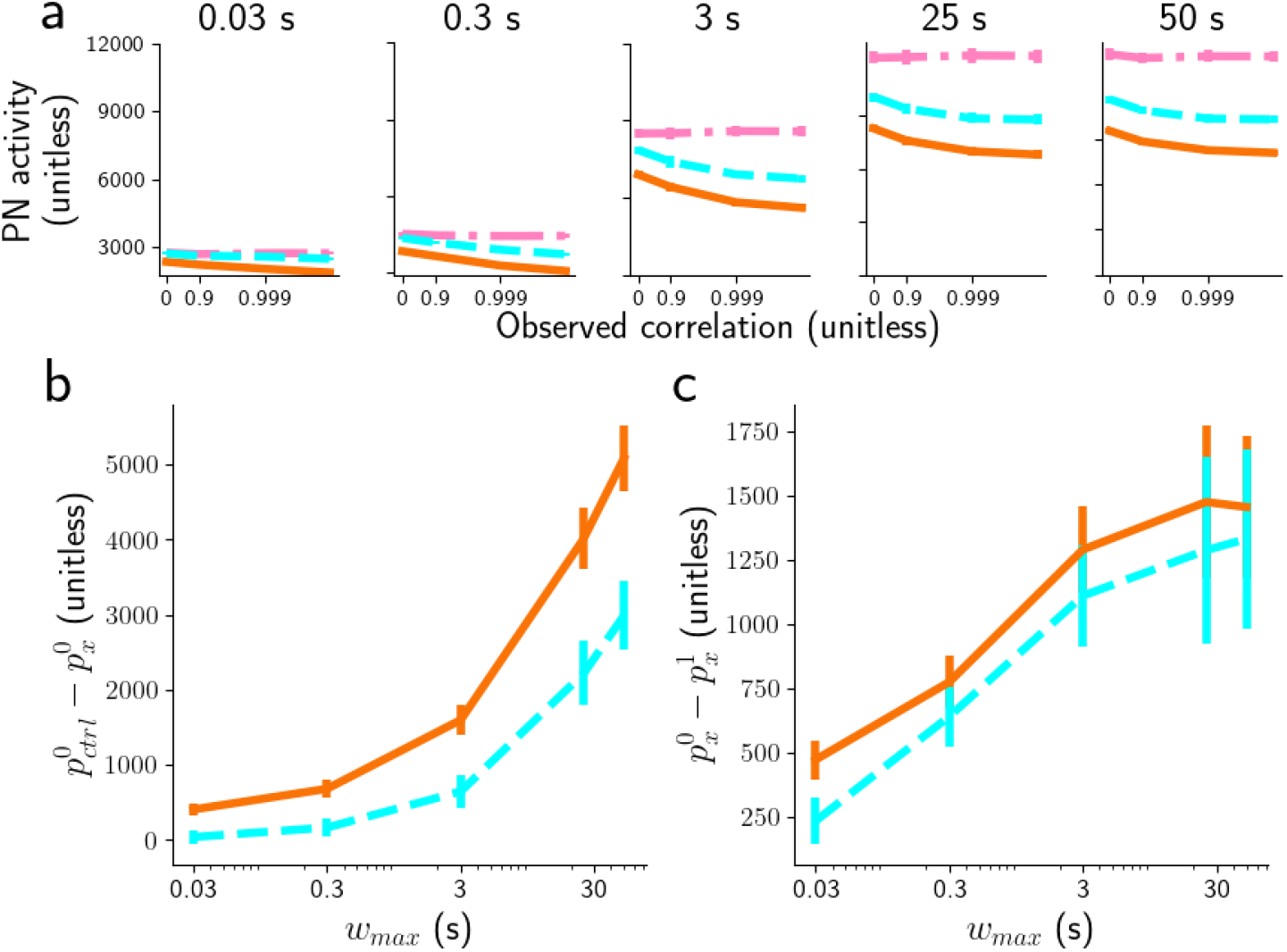
Similar results Fig.9 using threshold 50 Hz. a) peak PN threshold 50 Hz for different subsets of whiff durations (from 0.01 to 50 s) for the three models: control model (dot dashed pink), LN model (orange continuous), and NSI model (dashed cyan). Note that the horizontal axis has a log-scale.

**Fig S15.**
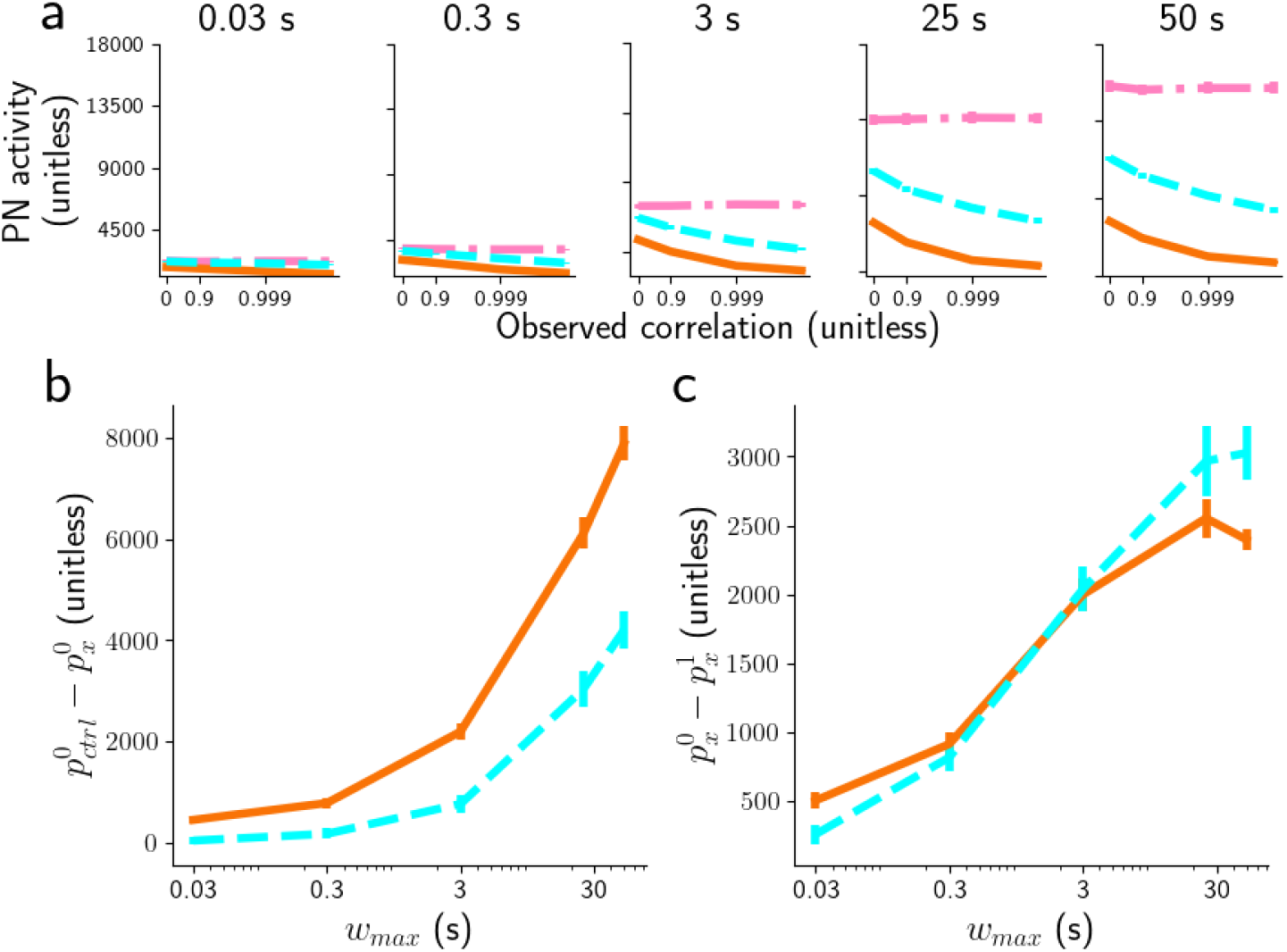
Similar results of Fig.9 using threshold 100 Hz. a) peak PN threshold 100 Hz for different subsets of whiff durations (from 0.01 to 50 s) for the three models: control model (dot dashed pink), LN model (orange continuous), and NSI model (dashed cyan). Note that the horizontal axis has a log-scale.

